# Impact of lossy compression of nanopore raw signal data on basecalling and consensus accuracy

**DOI:** 10.1101/2020.04.19.049262

**Authors:** Shubham Chandak, Kedar Tatwawadi, Srivatsan Sridhar, Tsachy Weissman

## Abstract

**Motivation:** Nanopore sequencing provides a real-time and portable solution to genomic sequencing, enabling better assembly, structural variant discovery and modified base detection than second generation technologies. The sequencing process generates a huge amount of data in the form of raw signal contained in fast5 files, which must be compressed to enable efficient storage and transfer. Since the raw data is inherently noisy, lossy compression has potential to significantly reduce space requirements without adversely impacting performance of downstream applications.

**Results:** We explore the use of lossy compression for nanopore raw data using two state-of-the-art lossy time-series compressors, and evaluate the tradeoff between compressed size and basecalling/consensus accuracy. We test several basecallers and consensus tools on a variety of datasets at varying depths of coverage, and conclude that lossy compression can provide 35-50% further reduction in compressed size of raw data over the state-of-the-art lossless compressor with negligible impact on basecalling accuracy (≲0.2% reduction) and consensus accuracy (≲0.002% reduction). In addition, we evaluate the impact of lossy compression on methylation calling accuracy and observe that this impact is minimal for similar reductions in compressed size, although further evaluation with improved benchmark datasets is required for reaching a definite conclusion. The results suggest the possibility of using lossy compression, potentially on the nanopore sequencing device itself, to achieve significant reductions in storage and transmission costs while preserving the accuracy of downstream applications.

**Availability:** The code is available at https://github.com/shubhamchandak94/lossy_compression_evaluation.

**Supplementary information:** Supplementary data are available at *Bioinformatics* online.

## 1 Introduction

Nanopore sequencing technologies developed over the past decade provide a real-time and portable sequencing platform capable of producing long reads, with important applications in completing genome assemblies and discovering structural variants associated with several diseases (Jain *et al.*, 2016). Nanopore sequencing consists of a membrane with pores where DNA passes through the pore leading to variations in current passing through the pore. This electrical current signal is sampled to generate the raw signal data for the nanopore sequencer and is then basecalled to produce the read sequence. Due to the continuous nature of the raw signal and high sampling rate, the raw signal data requires large amounts of space for storage, e.g., a typical 30x depth human sequencing experiment can produce terabytes of raw signal data, which is an order of magnitude more than the space required for storing the basecalled reads (Jain *et al.*, 2018).

Due to the ongoing research into improving basecalling technologies and the scope for further improvement in accuracy, the raw data needs to be retained to allow repeated analysis of the sequencing data. This makes compression of the raw signal data crucial for efficient storage and transport. There have been a couple of lossless compressors designed for nanopore raw signal data, namely, Picopore (Gigante, 2017) and VBZ (https://github.com/nanoporetech/vbz_compression/). Picopore simply applies gzip compression to the raw data, while VBZ, which is the current state-of-the-art tool, uses variable byte integer encoding followed by zstd compression. Although VBZ reduces the size of the raw data by 60%, the compressed size is still quite significant and further reduction is desirable. However, obtaining further improvements in lossless compression is challenging due to the inherently noisy nature of the current measurements.

In this context, lossy compression is a natural candidate to provide a boost in compression at the cost of certain amount of distortion in the raw signal. There have been several works on lossy compression for time series data, including SZ (Liang *et al.*, 2018) and LFZip (Chandak *et al.*, 2020) that provide a guarantee that the reconstruction lies within a certain user-defined interval of the original value for all time steps. However, in the case of nanopore raw current signal, the metric of interest is not the maximum deviation from the original value, but instead the impact on the performance of basecalling and other downstream analysis steps. In particular, two quantities of interest are the basecalling accuracy and the consensus accuracy. The basecalling accuracy measures the similarity of the basecalled read sequence to the known true genome sequence, while the consensus accuracy measures the similarity of the consensus sequence obtained from multiple overlapping reads to the known true genome sequence. As discussed in Wick *et al.* (2019), these two measures are generally correlated but can follow different trends in the presence of systematic errors. In general, consensus accuracy can be thought of as the higher-level metric, which is usually of interest in most applications, while basecalling accuracy is a lower-level metric in the sequencing analysis pipeline.

In this work, we study the impact of lossy compression of nanopore raw signal data on basecalling and consensus accuracy. We evaluate the results for several basecallers and at multiple stages of the consensus pipeline to ensure the results are generalizable to future iterations of these tools. We find that lossy compression using general-purpose tools can provide significant reduction in file sizes with negligible impact on accuracy. To further stress-test the ability of lossy compression to preserve useful information, we look into the impact of lossy compression on methylation calling performance and reach similar conclusions as those for basecalling accuracy. To the best of our knowledge, this is the first study exploring the use of lossy compression for nanopore raw signal data and performing a systematic analysis of its impact on downstream applications. We believe our results provide motivation for research into specialized lossy compressors for nanopore raw signal data and suggest the possibility of reducing the resolution of the raw signal generated on the nanopore device itself while preserving the downstream performance. The source code and data for our analysis is publicly available at https://github.com/shubhamchandak94/lossy_compression_evaluation and can be useful as a benchmarking pipeline for further research into lossy compression for nanopore data.

## 2 Background

### 2.1 Nanopore sequencing and basecalling

Nanopore sequencing, specifically the MinION sequencer developed by Oxford Nanopore Technologies (ONT) (Jain *et al.*, 2016), involves a strand of DNA passing through a pore in a membrane with a potential applied across it. Depending on the sequence of bases present in the pore (roughly 6 bases at any instant), the ionic current passing through the pore varies with time and is measured at a sampling frequency of 4 kHz. The sequencing produces 5-15 current samples per base, which are quantized to a 16-bit integer and stored as an array in a version of the HDF5 format called fast5. The current signal is then processed by the basecaller to generate the basecalled read sequence and the corresponding quality value information. In the uncompressed format, the raw current signal requires close to 18 bytes per sequenced base which is significantly more than the amount needed for storing the sequenced base and the associated quality value. The sequenced FASTQ files can also be compressed further using specialized compressors (Dufort y Álvarez *et al.*, 2020) to further reduce the storage costs.

Over the past years, there has been a shift in the basecalling strategy from a physical model-based approach to a machine learning-based approach leading to significant improvement in basecalling accuracy (see Rang *et al.* (2018) and Wick *et al.* (2019) for a detailed review). In particular, the current default basecaller Guppy by ONT (based on open source tool Flappie (https://github.com/nanoporetech/flappie)) uses a recurrent neural network that generates transition probabilities for the bases which are then converted to the most probable sequence of bases using Viterbi algorithm. Another recent basecaller by ONT is bonito (https://github.com/nanoporetech/bonito/, currently experimental), which is based on a convolutional neural network and CTC decoding (Graves *et al.*, 2006), achieving close to 92-95% basecalling accuracy in terms of edit distance. Despite the progress in basecalling, the current error rates are still relatively high (typically 5-10%) with considerable fraction of insertion and deletion errors, which necessitates the storage of the raw data for utilizing improvements in the basecalling technologies for future (re)analysis.

### 2.2 Assembly, consensus and polishing

Long nanopore reads allow much better repeat resolution and are able to capture long-range information about the genome leading to significant improvements in de novo genome assembly (Jain *et al.*, 2018). However genome assembly with nanopore data needs to handle the much higher error rates as compared to second generation technologies such as Illumina, and there have been several specialized assemblers for this purpose, including Flye (Kolmogorov *et al.*, 2019; Lin *et al.*, 2016) and Canu (Koren *et al.*, 2017), some of which allow hybrid assembly with a combination of short and long read data.

Nanopore de novo assembly is usually followed by a consensus step that improves the assembly quality by aligning the reads to a draft assembly and then performing consensus from overlapping reads (e.g., Racon (Vaser *et al.*, 2017)). Note that the consensus step can be performed even without de novo assembly if a reference sequence for the species is already available, in which case the alignment to the reference is used to determine the overlap between reads. Further polishing of the consensus sequence can be performed with tools specialized for nanopore sequencing that use the noise characteristics of the sequencing and/or basecalling to find the most probable consensus sequence. For example, Nanopolish (Loman *et al.*, 2015) directly uses the raw signal data for polishing the consensus using a probabilistic model for the generation of the raw signal from the genomic sequence. Medaka (https://nanoporetech.github.io/medaka/) is the current state-of-the-art consensus polishing tool both in terms of runtime and accuracy (https://github.com/rrwick/August-2019-consensus-accuracy-update/). Medaka uses a neural network to perform the consensus from the pileup of the basecalled reads at each position of the genome.

### 2.3 Methylation calling

DNA methylation plays an important role in various biological functions (Simpson *et al.*, 2017), with 6mA and 5mC being the most commonly studied methylated bases (methylated versions of A and C, respectively). Since nanopore sequencing can work with native (non-PCR amplified) DNA, it is possible to detect methylated bases due to the changes in the raw signal when the methylated base passes through the pore. This fact has been exploited to develop methylation calling pipelines using various techniques including hidden Markov model (HMM)-based methods and neural network-based methods (Loman *et al.*, 2015; Simpson *et al.*, 2017; Liu *et al.*, 2019; Ni *et al.*, 2019). In this work, we use Megalodon (https://github.com/nanoporetech/megalodon/) which first anchors the intermediate probabilities produced by the basecalling neural network (from Guppy basecalling modes that call both modified and unmodified bases) to the reference sequence. It then uses traditional HMM algorithms such as Viterbi and forward-backward algorithms (Rabiner, 1989) to compute the probability that a given base is modified. This common framework can be used to call different types of base modifications by providing the appropriate basecalling model, and we focus on CpG motifs on the human genome in this work.

### 2.4 Lossy compression

Lossy compression (Gersho and Gray, 2012) refers to compression of the data into a compressed bitstream where the decompressed (reconstructed) data need not be exactly but only approximately similar to the original data. Lossy compression is usually studied in the context of a distortion metric that specifies how the distortion or error between the original and the reconstruction is measured. This gives rise to a tradeoff between the compressed size and the distortion, referred to as the rate-distortion curve. Here, we work with two state-of-the-art lossy compressors for time-series data, LFZip (Chandak *et al.*, 2020) and SZ (Liang *et al.*, 2018). Both these compressors work with a maxerror parameter that specifies the maximum absolute deviation between the original and the reconstructed data. If *x*_1_,…, *x*_*T*_ denotes the original time-series, 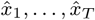 denotes the reconstructed time-series, and *E* is the maxerror parameter, then these compressors guarantee that 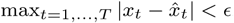.

LFZip and SZ use slightly different approaches towards lossy compression. LFZip uses a prediction-quantization-entropy coding approach, and in this work we use a mode wherein the prediction step is skipped and LFZip simply performs uniform scalar quantization (i.e., uniform binning with the bin size determined by the maxerror) followed by entropy coding. This mode provided the best compression in our experiments. SZ uses a curve fitting step followed by entropy coding, with the reconstruction lying on a low-degree local polynomial approximation to the original data.

There are a couple of reasons for focusing on lossy compressors with maximum absolute deviation as the distortion metric instead of mean square error or mean absolute error in this work. The first reason is that the guaranteed maximum error implies that the reconstructed raw signal is close to the original value at each and every timestep and not only in the average sense. Hence, the maximum error distortion metric is preferable for general applications where the true distortion metric is not well understood. The second reason is the availability of efficient implementations which is crucial for compressing the large genomic datasets. However, we believe that there is significant scope for using mean square error and other metrics for developing specialized lossy compressors for nanopore data given the better theoretical understanding of those metrics.

## 3 Experiments

We next describe the experimental setup in detail (see Figure 1 for a flowchart representation). The instructions for downloading the datasets and installing the tools, as well as the scripts for performing the experiments are available on the GitHub repository. The experiments were run on an Ubuntu 18.04.4 server with 40 Intel Xeon processors (2.2 GHz), 260 GB RAM and 8 Nvidia TITAN X (Pascal) GPUs.

**Fig. 1.**
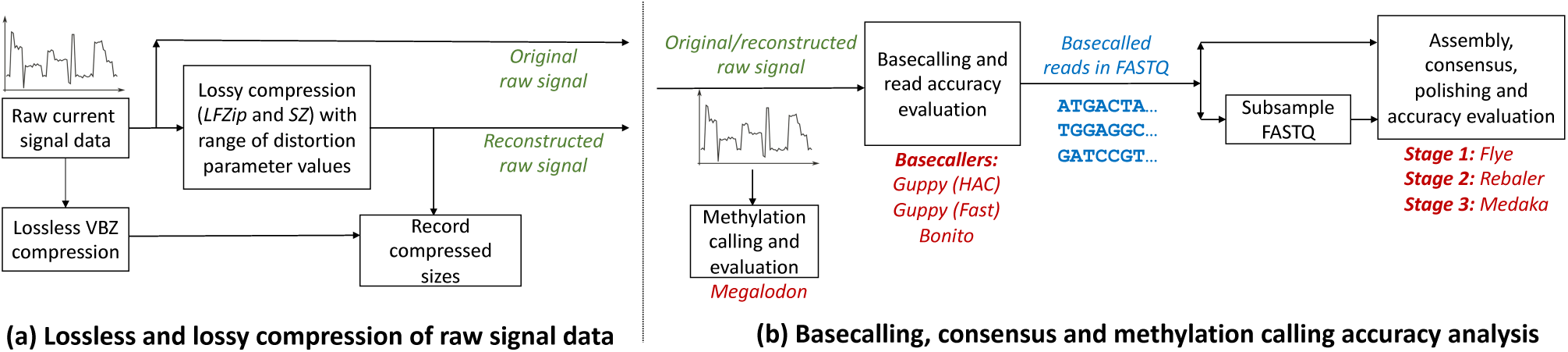
Flowchart showing the experimental procedure. (a) The raw data was compressed with both lossless and lossy compression tools, (b) the original and lossily compressed data was then basecalled with three basecalling tools. Finally, the basecalled data and its subsampled versions were assembled and the assembly (consensus) was polished using a three-step pipeline (1. Flye, 2. Rebaler, 3. Medaka). The tradeoff between compressed size and basecalling/consensus accuracy was studied. For one dataset, methylation calling and accuracy evaluation was performed using Megalodon. Parts of the evaluation pipeline are based on previous work on basecaller comparison in Wick et al. (2019) and its addendum (https://github.com/rrwick/August-2019-consensus-accuracy-update/).

### 3.1 Datasets

Table 1 shows the datasets used for analysis in this work. The first three bacterial datasets were chosen to be representative datasets with different GC-content and flowcell types, including the latest R10.3 pore. We note that some of the tools were not run on the *E. coli* dataset since they do not yet support the R10.3 pore. For all these datasets the ground-truth genomic sequence is known through hybrid assembly with long and short read technologies. The table also shows the uncompressed and VBZ compressed sizes for the datasets, we observe that lossless compression can provide size reduction of roughly 60%. For each dataset, we run our analysis on both the original read depth as well as subsampled versions of the datasets (2x, 4x and 8x subsampling of Fastq files performed using Seqtk (https://github.com/lh3/seqtk/)). This helps us understand how the impact of lossy compression on consensus accuracy depends on read depth. The last dataset (human) is used for basecalling and methylation calling accuracy evaluation, and we use one flowcell (FAB45280) of NA12878 human nanopore data from Jain *et al.* (2018) consisting of around 900M sequenced bases. For basecalling accuracy evaluation, we generate the ground truth genome by applying the variants from GIAB (Zook *et al.*, 2016) to the reference genome. More details on the methylation calling experiments are provided in Section 3.5.

**Table 1.** Datasets used for analysis. The E. coli dataset was obtained from http://albertsenlab.org/we-ar10-3-pretty-close-now/. N50 is a statistical measure of average length of the reads (see Wick et al. (2019) for a precise definition). The uncompressed size column refers to storing the raw signal in the default representation using 16 bits/signal value. The first three datasets (bacterial) were used for basecalling and consensus accuracy evaluation, while the last dataset (low-coverage human dataset from a single flowcell) was used for basecalling and per-read methylation calling accuracy evaluation.

| Species | Sample | Genome size (bp) | GC-content | Flowcell type | Read count | Read length N50 (bp) | Approx. depth | Raw signal size (GB) |  | Source |
| --- | --- | --- | --- | --- | --- | --- | --- | --- | --- | --- |
|  |  |  |  |  |  |  |  | Uncompressed | VBZ (lossless) |  |
| *Staphylococcus aureus* | CAS38_02 | $2.9 \times 10^6$ | 32.8% | R9.4.1 | 11,047 | 24,666 | 83x | 4.86 | 2.02 | Wick et al. (2019) |
| *Klebsiella pneumoniae* | INF032 | $5.1 \times 10^6$ | 57.6% | R9.4 | 15,154 | 37,181 | 108x | 10.14 | 4.32 | Wick et al. (2019) |
| *Escherichia coli* | K-12 MG1655 | $4.6 \times 10^6$ | 50.8% | R10.3 | 92,000 | 7,431 | 128x | 12.09 | 5.14 | See Caption |
| *Homo sapiens* | NA12878 | $3.1 \times 10^9$ | 40.9% | R9.4 | 128,314 | 11,404 | 0.29x | 25.37 | 10.31 | Jain et al. (2018) |

We focus on using bacterial datasets rather than human datasets for consensus accuracy evaluation for a few reasons similar to those cited in Wick *et al.* (2019). Firstly, bacterial datasets typically have a more reliable ground truth allowing for more precise estimation of the impact of lossy compression. This is especially important for consensus accuracy which can be very high (≳99.9%), leading to a much greater uncertainty in the evaluation due to errors in the ground truth sequence. The smaller size for bacterial datasets also allows more extensive experimentation at higher coverage and across several parameters. Due to these reasons, previous studies have often relied on bacterial datasets for consensus/assembly accuracy evaluation, and used human datasets only for basecalling accuracy evaluation. (e.g., see the works Teng *et al.* (2018) and Zeng *et al.* 2020) on novel basecalling algorithms). In addition, lossy compression with maximum deviation constraint is typically local in nature, with LFZip in particular performing uniform scalar quantization independently at each time step. Thus, the size of the genome should not impact the analysis and the results should generalize to larger genomes. Further experimentation on larger eukaryotic datasets remains part of future work as better benchmark datasets are obtained.

We also looked into the possibility of using data from Zymo microbial community standard (Nicholls *et al.*, 2019) which has been used to evaluate basecallers and consensus tools. However, we decided to use the previously described datasets for a couple of reasons. Firstly, the Zymo dataset is a metagenomic dataset and obtaining data for individual species requires additional analysis and introduces a possibility of erroneous conclusions. Secondly, several neural network models in the downstream pipeline (e.g., basecallers and Medaka consensus) are commonly trained on parts of the Zymo dataset, with different tools using different training and testing genomes. This makes the data unsuitable when comparing several tools due to overfitting concerns.

### 3.2 Lossy compression

To study the impact of lossy compression, we generate new fast5 datasets by replacing the raw signal in the original fast5 files with the reconstruction produced by lossy compression. We use open source general-purpose time-series compressors LFZip (Chandak *et al.*, 2020) (version 1.1) and SZ (Liang *et al.*, 2018) (version 2.1.8.3). Both the tools require a parameter representing the maximum absolute deviation (maxerror) of the reconstruction from the original. We conducted ten experiments for each tool by setting the maxerror parameter to 1, 2,…, 10. To put this in context, note that for a typical current range of 60 pA and the typical noise value of 1-2 pA, the maxerror settings of 1 and 10 correspond to current values of 0.17 pA and 1.7 pA, respectively (https://github.com/nanoporetech/kmer_models/). Finally, we note that both LFZip and SZ can compress millions of timesteps per second and hence can be used to compress the nanopore raw signal data in real time as it is produced by the sequencer.

### 3.3 Basecalling and consensus

We perform basecalling on the raw signal data (both original and lossily compressed) using two modes of Guppy (version 3.6.1) as well as with bonito (version 0.2.0, note that bonito is currently an experimental release). For Guppy, we use the default high accuracy mode (guppy_hac) and the fast mode (guppy_fast). Both the modes use the same general framework but differ in terms of the neural network architecture size and weights. We use these three basecaller settings to study whether lossy compression leads to loss of useful information that can be potentially exploited by future basecallers.

We use a three-step assembly, consensus and polishing pipeline based on the analysis and recommendations in https://github.com/rrwick/August-2019-consensus-accuracy-update/. The first step is de novo assembly using Flye (version 2.7.1) (Kolmogorov *et al.*, 2019; Lin *et al.*, 2016) which produces a basic draft assembly. The second step is consensus polishing of the Flye assembly using Rebaler (https://github.com/rrwick/Rebaler/, version v0.2.0) which runs multiple rounds of Racon (version 1.4.13) to produce a high quality consensus of the reads. Finally, the third step uses Medaka (version 1.0.3) by ONT that performs further polishing of the Rebaler consensus using a neural network-based approach. Note that the neural network model for Medaka needs to be chosen corresponding to the basecaller.

### 3.4 Evaluation metrics

For evaluating the basecalling and consensus accuracy, we use the pipeline presented for the task of basecaller comparison in Wick *et al.* (2019) and its addendum (https://github.com/rrwick/August-2019-consensus-accuracy-update/). The basecalled reads were aligned to the true genome sequence using minimap2 (Li, 2018) and the read’s basecalled identity was defined as the number of matching bases in the alignment divided by the total alignment length. We report only the median identity across reads in the results section (see Supplementary Material for details on accessing the per-read results). The consensus accuracy after each stage is computed in a similar manner, where instead of aligning the reads, we split the assembly into 10 kbp pieces and then find median identity across these pieces. Finally, we compute the basecalling and consensus Qscore using the Phred scale as Qscore = −10 log_10_(1 − identity) where the identity is represented as a fraction. We refer the reader to Wick *et al.* (2019) for further discussion on these metrics. In addition, we evaluate the accuracy of homopolymer sequences in the consensus using the fastmer.py script (https://github.com/jts/assembly_accuracy), since homopolymer calling has been identified as one of the main challenges of nanopore sequencing (Rang *et al.*, 2018).

### 3.5 Methylation calling and evaluation

We consider the impact of lossy compression on methylation calling to understand whether the loss in information due to compression leads to further degradation of methylation calling performance as compared to basecalling, given that methylation calling is a more sensitive task than basecalling. For evaluating this impact, we use the pipeline and benchmark dataset used in two previous works, DeepMod (Liu *et al.*, 2019) and DeepSignal (Ni *et al.*, 2019). We use NA12878 human nanopore data from Jain *et al.* (2018) which used native (non-PCR amplified) DNA and use a benchmark obtained from bisulfite sequencing from the ENCODE project (ENCFF835NTC) (ENCODE-Project-Consortium *et al.*, 2012). Following the procedure in (Ni *et al.*, 2019), we identify the high confidence positive and negative sites on the genome by restricting ourselves to sites with coverage at least five, and 100% positive or negative calls on both strands in the bisulfite dataset. This resulted in roughly 5.4M positive and 4.7M negative high confidence sites on the genome.

We then used Megalodon (version 2.1.0) on the nanopore dataset to obtain a list of per-read methylation calls (predicted probabilities) for each CpG motif in the read. We used a basecalling model specially trained for calling CpG methylation released in the Rerio repository (https://github.com/nanoporetech/rerio/). We then computed the precision, recall and AUC (area under ROC curve), restricting ourselves to the high confidence sites determined above. For the precision and recall computation, we used a threshold of 0.5 for the predicted methylation probabilities.

As shown in Table 1, we used only one flowcell of data consisting of around 900M sequenced bases. For this study, we focused exclusively on per-read evaluation and did not attempt to compute correlation of the methylation frequencies with the bisulfite data as done in Ni *et al.* (2019). We believe that the per-read evaluation should be indicative of the extent to which lossy compression leads to loss of information regarding methylation. We also found other datasets in the previous works (Simpson *et al.*, 2017; Liu *et al.*, 2019; Ni *et al.*, 2019) where ground truth positive and negative datasets were generated using methyltransferase enzyme and PCR-amplification, respectively. However, these datasets were generated with older pores (R7 or 2D technology) which are not supported by the modern methylation calling tools. Better benchmark datasets in the future can be helpful for more extensive evaluation.

## 4 Results and discussion

We now discuss the main results obtained from the experiments described above. Throughout the results and discussion, the compressed sizes for lossy compression are shown relative to the compressed size for VBZ lossless compression, where the lossless compression sizes are shown in Table 1. Additional results and plots are available in the Supplementary Material.

Figure 2 shows the variation of the size of the lossily compressed dataset with the maxerror parameter. We see that LFZip generally provides better compression than SZ at the same maxerror value, although this fact by itself does not guarantee a better tradeoff for the metrics of interest. We also see that lossy compression can provide significant size reduction over lossless compression even with relatively small maxerror (recall that maxerror of 1 in the 16-bit representation of the raw signal corresponds to 0.17 pA error in the current value). For example, at maxerror of 5, lossy compression can provide size reduction of around 50% over lossless compression and size reduction of around 70% over the uncompressed 16-bit representation.

**Fig. 2.**
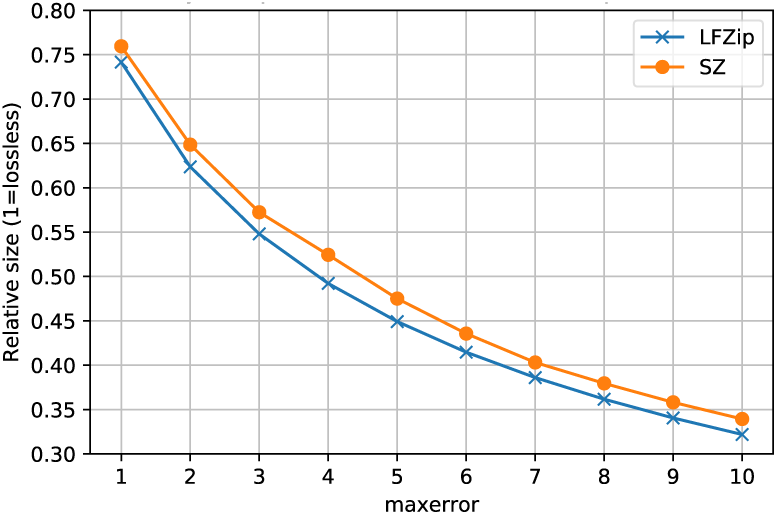
Compressed size for lossy compression with LFZip and SZ for the S. aureus dataset as a function of the maxerror parameter. The compressed sizes are shown relative to the VBZ lossless compression size.

Both SZ and LFZip can compress millions of samples per second, with SZ being about an order of magnitude faster than LFZip (Chandak *et al.*, 2020). Since LFZip simply performs uniform scalar quantization and entropy coding (in the mode used here), we believe that it can be significantly optimized further for this application. We also looked into the possibility that lossy compression could impact the speed/peak memory usage of the steps in the downstream pipeline (basecalling, assembly, etc.), however we observed that such effects were relatively small (≲5-10% change) and mostly attributable to experimental variation. See the Supplementary Material for time and memory usage results for various stages in the pipeline and the impact of lossy compression on these.

### Basecalling accuracy

Figure 3 shows the tradeoff achieved between basecalling accuracy and compressed size for the four datasets for all three basecallers. First, we observe that guppy_hac performs the best closely followed by bonito, and guppy_fast is typically far behind. As the maxerror parameter is increased, the basecalling accuracy stays stable for all the basecallers till the compressed size reaches 65% of the losslessly compressed size. For example, the basecalling accuracy for the *S. aureus* dataset with guppy_hac is *∼*96.1% for lossless compression, and *∼*96.0% at a 35% size reduction. After this the basecalling accuracy drops more sharply, becoming 2% lower than the original lossless level when the maxerror parameter is 10. The drop seems to follow a similar trend for all the basecallers and compressors, suggesting that at least 35% reduction in size over lossless compression can be obtained without sacrificing basecalling accuracy. Note that for maxerror parameter equal to 10, the allowed deviation of the reconstructed raw signal is larger than the typical noise levels in sequencing, and hence lossy compression probably leads to perceptible loss in the useful information contained in the raw signal.

**Fig. 3.**
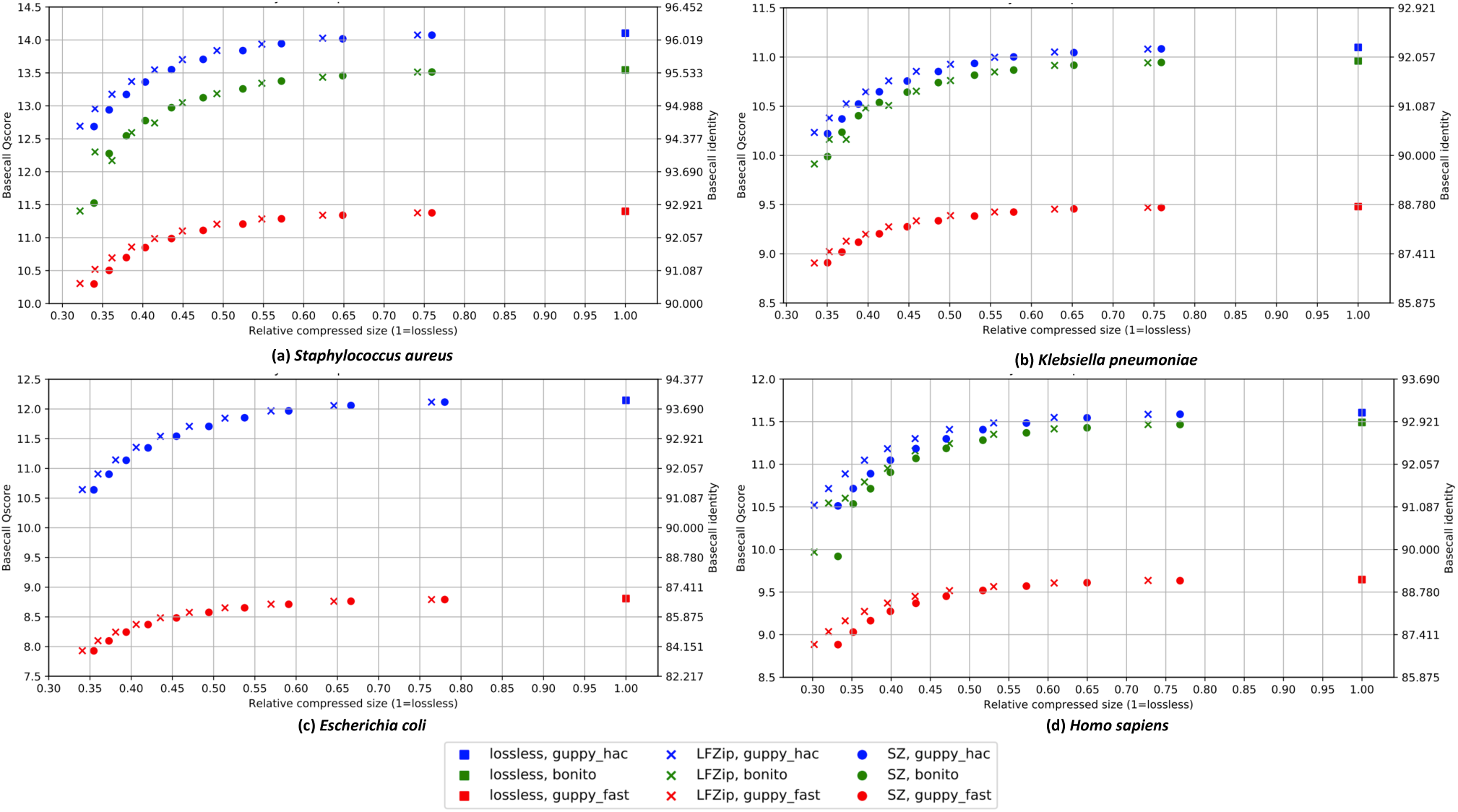
Basecalling accuracy vs. compressed size for (a) S. aureus, (b) K. pneumoniae, (c) E. coli, and (d) H. sapiens datasets. The results are displayed for the losslessly compressed data and the lossily compressed versions with LFZip and SZ (with maxerror 1 to 10) for the four basecallers. The compressed sizes are shown relative to the VBZ lossless compression size. Bonito was not run on E. coli due to lack of support for the R10.3 pore.

We observed that the impact of lossy compression on read lengths and the number of aligned reads is negligible (Supplementary Material). This suggests that lossy compression generally leads to local and small perturbations in the basecalled read and does not lead to major structural changes in the read such as loss of information due to trimmed/shortened reads. This is expected given that the lossy compressors used here guarantee that the reconstructed signal is within a certain deviation from the original signal at each time step.

### Consensus accuracy

Figures 4, 5(a) and 5(b) study the tradeoff between consensus accuracy and compressed size (i) across basecallers for the final Medaka polished assembly, (ii) across the assembly stages and (iii) across read depths in subsampled datasets, respectively. As expected, we observe that the consensus accuracy is significantly higher than basecalling accuracy across these experiments. We also observe that the consensus accuracy stays at the original lossless level till the compressed size reaches around 40-50% of the losslessly compressed size (50-60% reduction) and the drop in accuracy beyond this is relatively small. For example, the consensus accuracy for the *S. aureus* dataset with guppy_hac is *∼*99.997% for lossless compression, and stays the same at a 50% size reduction. Thus, the impact of lossy compression on consensus accuracy is less severe than that on basecalling accuracy. This suggests that the errors introduced by lossy compression are generally random in nature and are mostly corrected by the consensus process.

**Fig. 4.**
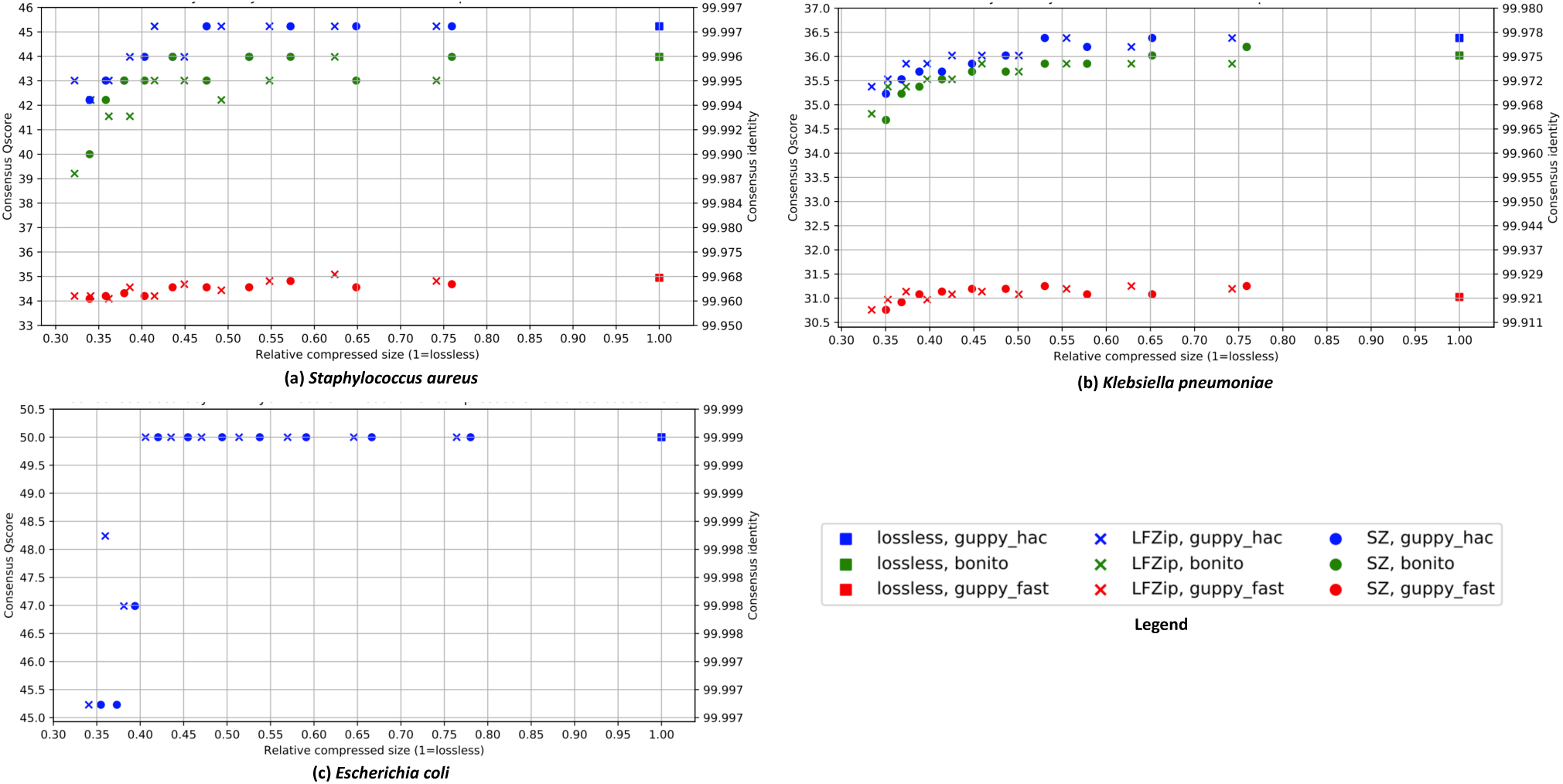
Consensus accuracy vs. compressed size for (a) S. aureus, (b) K. pneumoniae and (c) E. coli datasets. The results are displayed for the polished Medaka assembly for the losslessly compressed data and the lossily compressed versions with LFZip and SZ (with maxerror 1 to 10) for the four basecallers. The compressed sizes are shown relative to the VBZ lossless compression size. Bonito and guppy_fast were not used on E. coli due to lack of corresponding Medaka models for the R10.3 pore.

**Fig. 5.**
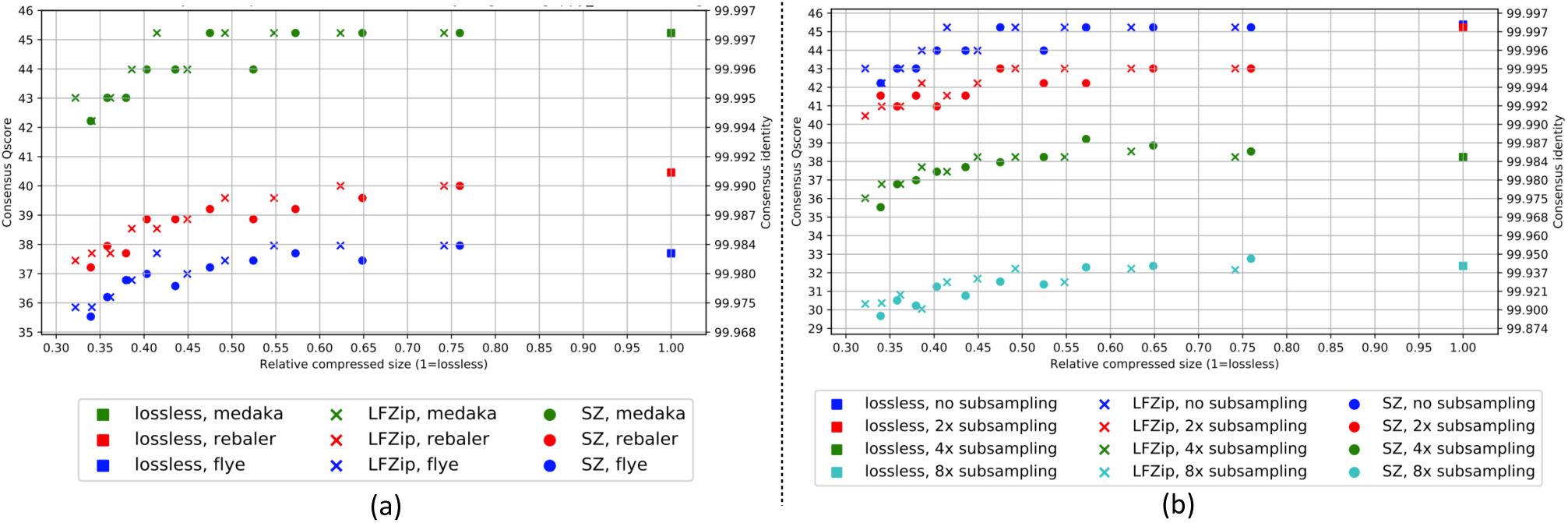
(a) Consensus accuracy vs. compressed size after each assembly step (Flye, Rebaler, Medaka) for the S. aureus dataset basecalled with guppy_hac. (b) Consensus accuracy (Medaka polished) vs. compressed size for subsampled versions (original, 2X subsampled, 4X subsampled, 8X subsampled) of the S. aureus dataset basecalled with guppy_hac. The results are displayed for the losslessly compressed data and the lossily compressed versions with LFZip and SZ (with maxerror 1 to 10). The compressed sizes are shown relative to the VBZ lossless compression size.

In our experiments, lossy compression did not affect the number of assembled contigs (always 1) and the contig length in most cases (Supplementary Material). The only exceptions were the 4x and 8x subsampled versions of the *E. coli* dataset where the assemblies for both the lossless and the lossily compressed datasets were fragmented. This might be due to lower data quality as evidenced by the significantly smaller read lengths for this dataset (see Table 1). In general, this again suggests that the impact of lossy compression is localized and without large-scale disruptions in the assembly/consensus process, although further experiments on larger eukaryotic genomes might be required to strengthen this claim.

Figure 5(a) considers the consensus accuracy after each stage of the assembly/consensus process (Flye, Rebaler, Medaka) for the *S. aureus* dataset basecalled with guppy_hac. We see that each stage leads to further improvement in the consensus accuracy. We also observe that the earlier stages of the pipeline are impacted more heavily by lossy compression (in terms of percentage reduction in accuracy) than the final Medaka stage. This is expected since each successive stage of the assembly/consensus pipeline provides further correction of the basecalling errors caused due to lossy compression. This effect is similar to the equalizing effect of polishing applied to different basecallers observed in Wick *et al.* (2019). We see a similar trend for the other dataset and analysis tools (see Supplementary Material).

Figure 5(b) studies the impact of subsampling to lower read depths on the consensus accuracy (after Medaka polishing) for the *S. aureus* dataset basecalled with guppy_hac. Note that the original dataset has around 80x depth of coverage, so 8x subsampling produces a depth of 10x which is generally considered quite low. We observe that lossy compression has more severe impact on consensus accuracy for lower depths, but 40-50% of size reduction can still be achieved without sacrificing the accuracy. This is again expected because consensus works better with higher depth datasets and is able to correct a greater fraction of the basecalling errors. We see a similar trend for the other dataset and analysis tools (see Supplementary Material).

Figure 6 considers the accuracy of homopolymers (of length 5 to 8) for the Medaka polished assembly of the *S. aureus* dataset basecalled with guppy_hac. We see that the impact of lossy compression is more pronounced for longer homopolymer sequences which are harder for the basecaller and assembly tools to handle, with around 30% size reduction over lossless compression possible with negligible impact on the accuracy. Thus, depending on the requirements, a lower maxerror parameter should be chosen to achieve higher accuracy for the longer homopolymer sequences. We believe it should be possible to overcome this challenge by designing specialized lossy compressors and by training the models in the basecalling and consensus pipeline on the lossily compressed data. We see a similar trend for other subsampling levels and other datasets (see Supplementary Material).

**Fig. 6.**
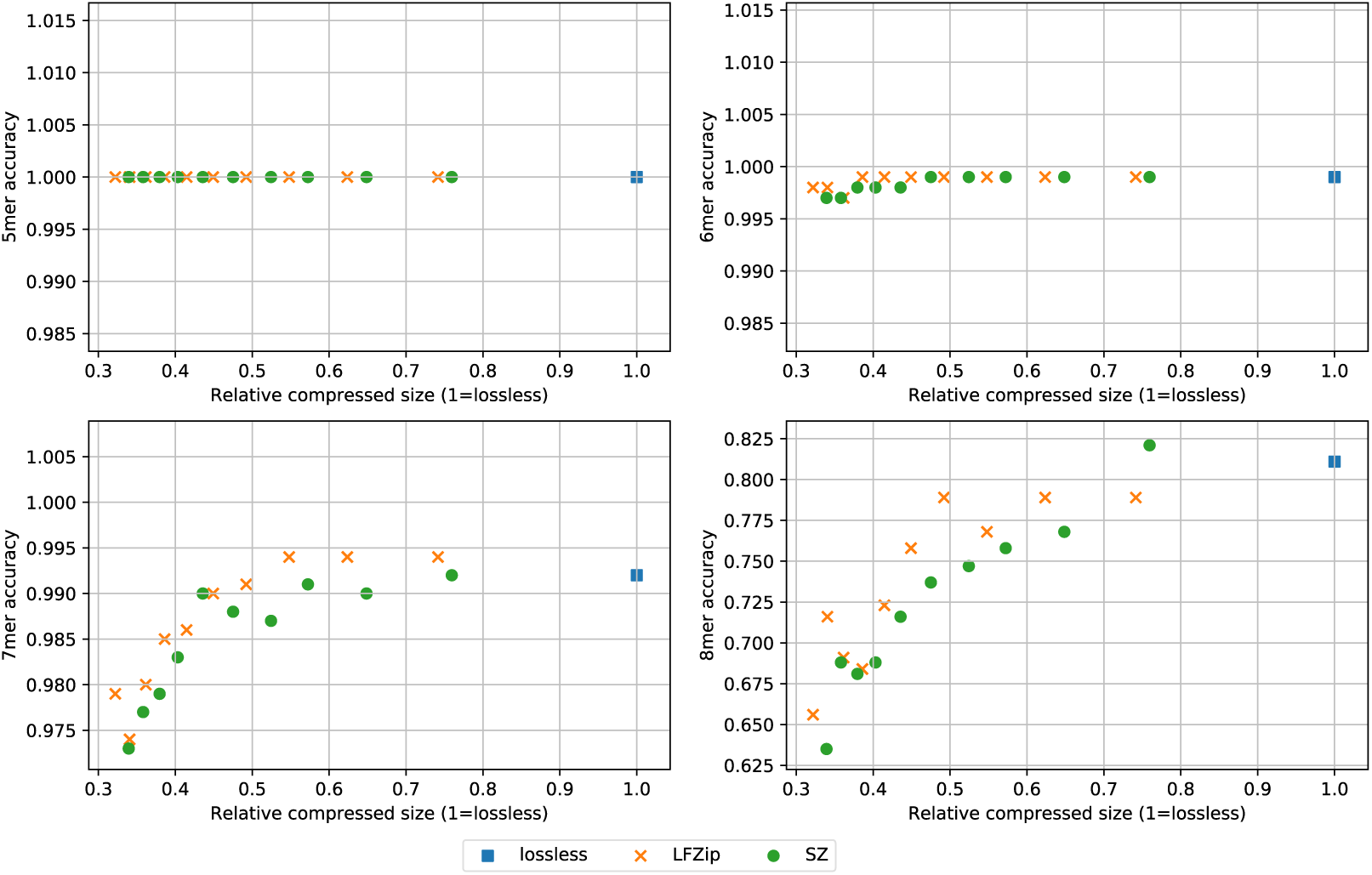
Consensus accuracy (Medaka polished) for homopolymer sequences of length 5 to 8 for the S. aureus dataset basecalled with guppy_hac. The results are displayed for the losslessly compressed data and the lossily compressed versions with LFZip and SZ (with maxerror 1 to 10). The compressed sizes are shown relative to the VBZ lossless compression size.

Overall, we observe that both LFZip and SZ can be used as tools to significantly save on space without sacrificing basecalling and consensus accuracy. The savings in space are close to 50% over lossless compression and 70% over the uncompressed representation. While it is not possible to say with certainty that we don’t lose any information in the raw signal that might be utilized by future basecallers, the results for the different basecallers and consensus stages suggest that applying lossy compression (for a certain range of parameters) only affects the noise in the raw signal without affecting the useful components. Finally, the decision to apply lossy compression and the extent of lossy compression should be based on the read depth (coverage), with more savings possible at higher depths where consensus accuracy is the metric of interest.

### Methylation calling accuracy

Figure 7 shows the precision, recall and AUC for CpG methylation calling on the NA12878 *H. sapiens* dataset. Across the >128K reads, there were roughly 670K positive ground truth positions and 560K negative ground truth positions after alignment. Recall that only genomic positions with sufficiently high confidence regarding the methylation status from bisulfite data were considered for the evaluation (*∼*5.4M positive and *∼*4.7M negative sites on genome, counting both strands). The achieved precision, recall and AUC were roughly 0.945, 0.845 and 0.945 respectively. As seen in the figure, the impact of lossy compression is similar to that on basecalling accuracy, with negligible impact for compression gains around 35-40% over lossless compression. This is despite the fact that methylation calling is generally considered a harder problem than basecalling due to the increased resolution needed for it. Further benchmarking using improved benchmark datasets in the future can be performed to strengthen these conclusions.

**Fig. 7.**
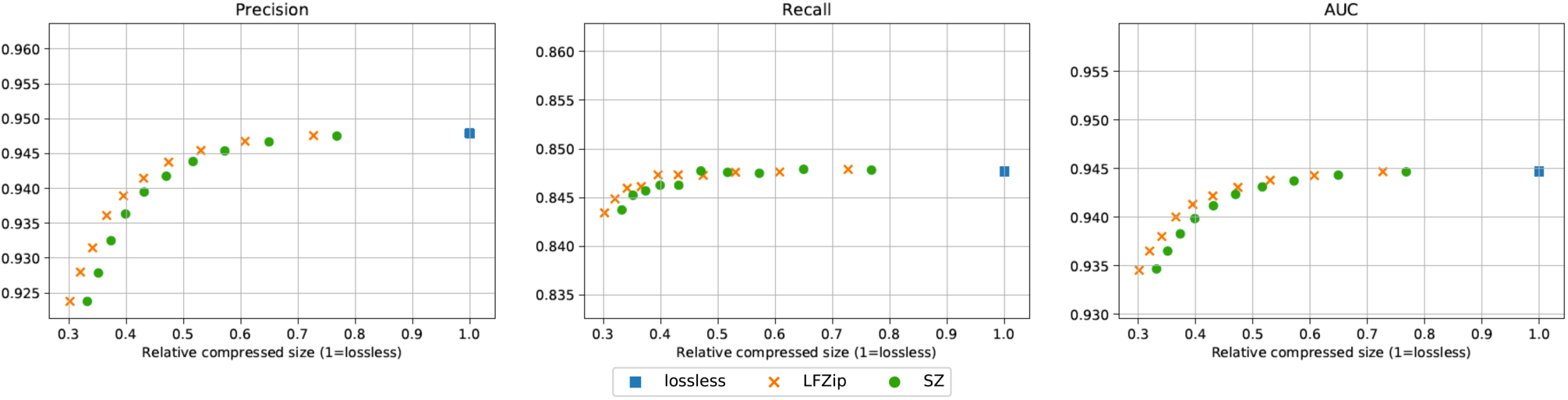
Precision, recall and AUC (area under ROC curve) for NA12878 CpG methylation calling using Megalodon. The metrics are computed for per-read methylation calls. For the precision and recall, a probability threshold of 0.5 was used for the predicted methylation probabilities. The results are displayed for the losslessly compressed data and the lossily compressed versions with LFZip and SZ (with maxerror 1 to 10). The compressed sizes are shown relative to the VBZ lossless compression size.

## 5 Conclusions and future work

We explore the use of lossy compression for nanopore raw data and its imapct on the basecalling and consensus accuracy. We find that lossy compression with existing tools can reduce the compressed size by 35-50% over lossless compression with less than 0.2% percent reduction in basecalling accuracy. The impact on consensus accuracy is even lower with less than 0.002% reduction at similar compression levels. Similar conclusions hold across datasets at different depths of coverage as well as several basecalling and assembly stages (with slight variation in the impact due to baseline lossless levels), suggesting that lossy compression with appropriate parameters does not lead to loss of useful information in the raw signal. For datasets with high depth of coverage, even further reduction is possible without sacrificing consensus accuracy. The analysis pipeline and data, partly based on Wick *et al.* (2019) and its addendum, are available online on GitHub along with documentation, and can be useful for further experimentation and development of specialized lossy compressors for nanopore raw signal data, which is part of future work. Further experiments on methylation accuracy evaluation and assembly for larger eukaryotic genomes are also part of future work, contingent upon the availability of improved benchmark datasets.

We believe that further research in this direction can lead to lossy compression algorithms tuned to the specific structure of the nanopore data and the evaluation metrics of interest, leading to further reduction in the compressed size. Further research into modeling the raw signal and the noise characteristics can help in this front. Another interesting direction could be the possibility of jointly designing the lossy compression with the modification of the algorithms in the downstream applications to match this compression. In particular, the current neural network models used in the basecallers can be retrained on the lossily compressed data to further understand the loss in information due to lossy compression. Finally, just as research on impact of lossy compression of Illumina quality scores on variant calling (Yu *et al.*, 2015; Ochoa *et al.*, 2017) led to Illumina reducing the default resolution of quality scores, it might be interesting to explore a similar possibility for nanopore data by performing the lossy compression or data binning on the nanopore sequencing device itself.

## Acknowledgements

We thank Pulkit Tandon for helpful discussions on LFZip.

