## Supplementary Material for "Impact of lossy compression of nanopore raw signal data on basecalling and consensus accuracy"

**Code and raw outputs of experiments available on GitHub at:**
`https://github.com/shubhamchandak94/lossy_compression_evaluation`

### 1 Time and memory usage

In this section, we provide the time and peak memory usage for lossy/lossless compression, as well as for the steps in the downstream analysis. We focus on the *S. aureus* dataset, with three settings of maxerror (1, 5, 10) for the lossy compressors, and basecalling done with the default basecaller Guppy (high accuracy mode). The three maxerror settings allow comparison across different levels of lossy compression. All experiments were performed on an Ubuntu 18.04.4 server with 40 Intel Xeon processors (2.2 GHz), 260 GB RAM and 8 Nvidia TITAN X (Pascal) GPUs. The default Python version was 3.7.6 (Anaconda).

| Mode | Time (s) | Peak memory (MB) | Average #samples/s |
|---|---|---|---|
| VBZ (lossless) | 2135 | 91.5 | 1.14M |
| LFZip (maxerror 1) | 4117 | 64.6 | 0.59M |
| SZ (maxerror 1) | 814 | 66.9 | 2.99M |
| LFZip (maxerror 5) | 3998 | 67.9 | 0.61M |
| SZ (maxerror 5) | 773 | 68.2 | 3.14M |
| LFZip (maxerror 10) | 3879 | 67.9 | 0.63M |
| SZ (maxerror 10) | 697 | 68.1 | 3.49M |

Table 1: Time and peak memory usage for compression+decompression of the *S. aureus* dataset for different compressors and maxerror parameters. The lossless compression time only includes the compression time. The last column shows the average number of raw signal samples handled per second, where the total number of raw signal samples for this dataset is roughly 2.43 billion.

Table 1 shows the time and memory usage for the compression stage. Due to the way the time was measured, for lossy compressors (LFZip, SZ) it includes the time for loading the reads from the original fast5, compressing the raw signal, decompressing, and writing the reconstructed signal to a new fast5. For lossless compressor VBZ, the time includes the time for loading the reads from the original fast5 and compressing the raw signal. Also note that we ran VBZ in the highest compression mode (22) since we aimed to compare with the state-of-the-art lossless compressor. The default application with lower compression mode (1) can be significantly faster. All compressors were run on the CPU with a single thread.

From Table 1, we see that generally SZ is 5-6 times faster than LFZip, and the compression speed slightly increases with higher maxerror settings. Note that the lossy compression speed is in the order of millions of raw signal samples per second, while the nanopore sampling frequency is 4000 samples/s for DNA pores. Thus, it should be possible to integrate lossy compression in a real-time manner with the sequencing process, especially upon improved integration with the pipeline and further optimization. We note that the peak memory usage is independent of the number of reads, and hence shouldn't be a factor in the scalability of these algorithms.

| Mode | Time (s) | Peak memory (GB) |
|---|---|---|
| VBZ (lossless) | 927 | 6.20 |
| LFZip (maxerror 1) | 969 | 7.42 |
| SZ (maxerror 1) | 952 | 6.67 |
| LFZip (maxerror 5) | 1005 | 6.68 |
| SZ (maxerror 5) | 997 | 6.78 |
| LFZip (maxerror 10) | 989 | 7.14 |
| SZ (maxerror 10) | 979 | 6.97 |

Table 2: Time and peak memory usage for Guppy (high accuracy) basecalling of the *S. aureus* dataset for different compressors and maxerror parameters.

Table 2 shows the time and memory usage for Guppy (high accuracy), which was run using a GPU. We see a very small increase in the time and memory usage for lossy compression as compared to lossless compression, but it is hard to distinguish from experimental noise and we can conclude that the impact of lossy compression the computational requirements for basecalling is relatively minor for this dataset.

| Mode | Time (s) | Peak memory (GB) |
|---|---|---|
| VBZ (lossless) | 1020 | 4.58 |
| LFZip (maxerror 1) | 1003 | 4.31 |
| SZ (maxerror 1) | 1054 | 5.12 |
| LFZip (maxerror 5) | 1007 | 4.49 |
| SZ (maxerror 5) | 993 | 4.53 |
| LFZip (maxerror 10) | 885 | 4.47 |
| SZ (maxerror 10) | 936 | 4.05 |

Table 3: Time and peak memory usage for Flye assembly of the *S. aureus* dataset for different compressors and maxerror parameters.

| Mode | Time (s) | Peak memory (GB) |
|---|---|---|
| VBZ (lossless) | 1088 | 1.04 |
| LFZip (maxerror 1) | 1100 | 1.01 |
| SZ (maxerror 1) | 1102 | 0.97 |
| LFZip (maxerror 5) | 1122 | 1.03 |
| SZ (maxerror 5) | 1111 | 1.04 |
| LFZip (maxerror 10) | 1178 | 1.23 |
| SZ (maxerror 10) | 1189 | 1.19 |

Table 4: Time and peak memory usage for Rebaler consensus of the *S. aureus* dataset for different compressors and maxerror parameters.

| Mode | Time (s) | Peak memory (GB) |
|---|---|---|
| VBZ (lossless) | 109 | 8.07 |
| LFZip (maxerror 1) | 93 | 8.13 |
| SZ (maxerror 1) | 93 | 8.82 |
| LFZip (maxerror 5) | 95 | 7.66 |
| SZ (maxerror 5) | 95 | 7.34 |
| LFZip (maxerror 10) | 98 | 9.95 |
| SZ (maxerror 10) | 98 | 9.11 |

Table 5: Time and peak memory usage for Medaka polishing of the *S. aureus* dataset for different compressors and maxerror parameters.

Tables 3, 4 and 5 show the time and memory usage for the three assembly/consensus stages: Flye, Rebaler and Medaka, respectively. All three tools were run using 8 CPU threads. We see a small variation in the time and memory usage for lossy compression as compared to lossless compression (particularly at high maxerror), but generally the variation is hard to distinguish from experimental noise and we can conclude that the impact of lossy compression the computational requirements for assembly/consensus is relatively minor.

#### 2 Supplementary plots

This section contains additional plots pertaining to different experiments that are not included in the main text for brevity. The Jupyter notebooks used for generating these plots are available on GitHub at `https://github.com/shubhamchandak94/lossy_compression_evaluation/tree/master/plots` in the respective subdirectories. The raw data for the plots is available in the form of tsv files at `https://github.com/shubhamchandak94/lossy_compression_evaluation/tree/master/data`.

- The results are displayed for the losslessly compressed data and the lossily compressed versions with LFZip and SZ (with maxerror 1 to 10). The compressed sizes are shown relative to the VBZ lossless compression size.
- Some tools were not run on the *E. coli* dataset due to lack of support for R10.3 pore.
- More details on the experiments and the plots can be found in the main paper and in the code on GitHub.

##### 2.1 *S. aureus*

###### 2.1.1 Lossy compression sizes

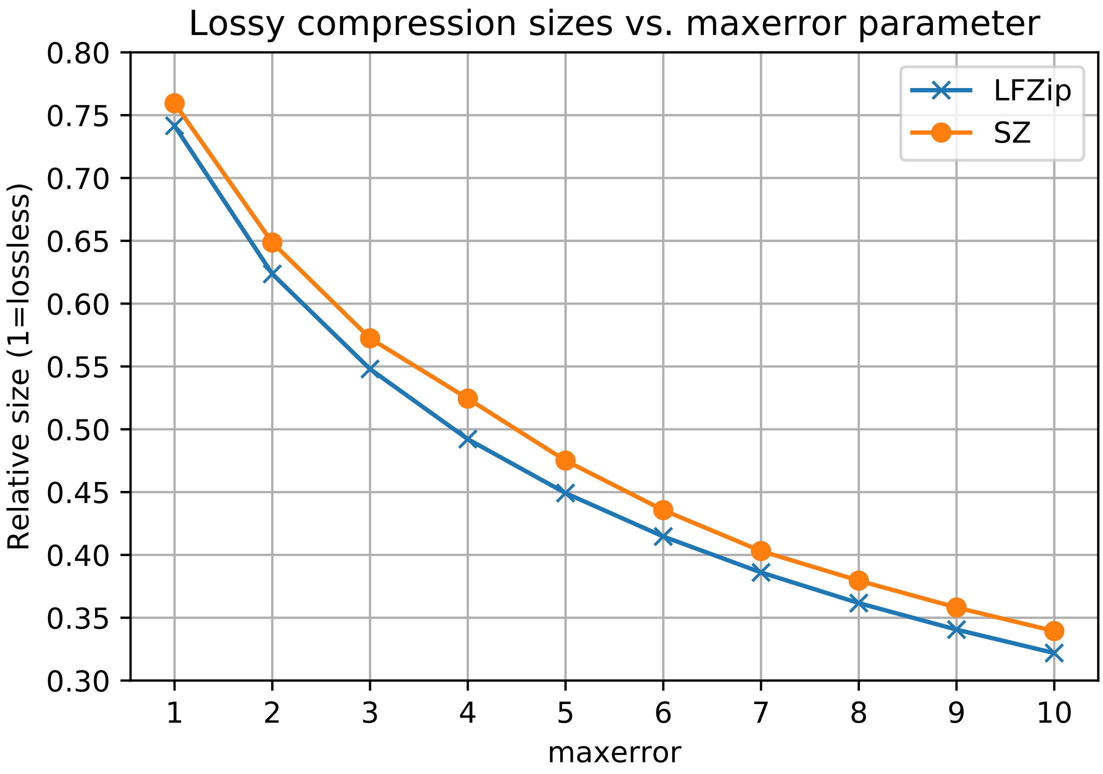

##### 2.1.2 Basecalling accuracy

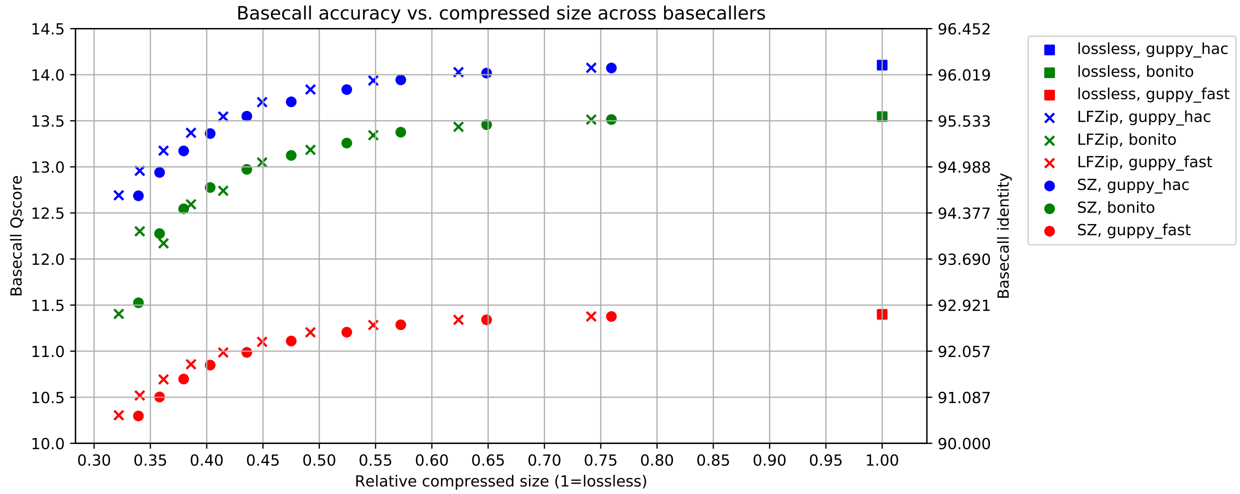

##### 2.1.3 Assembly accuracy across basecallers

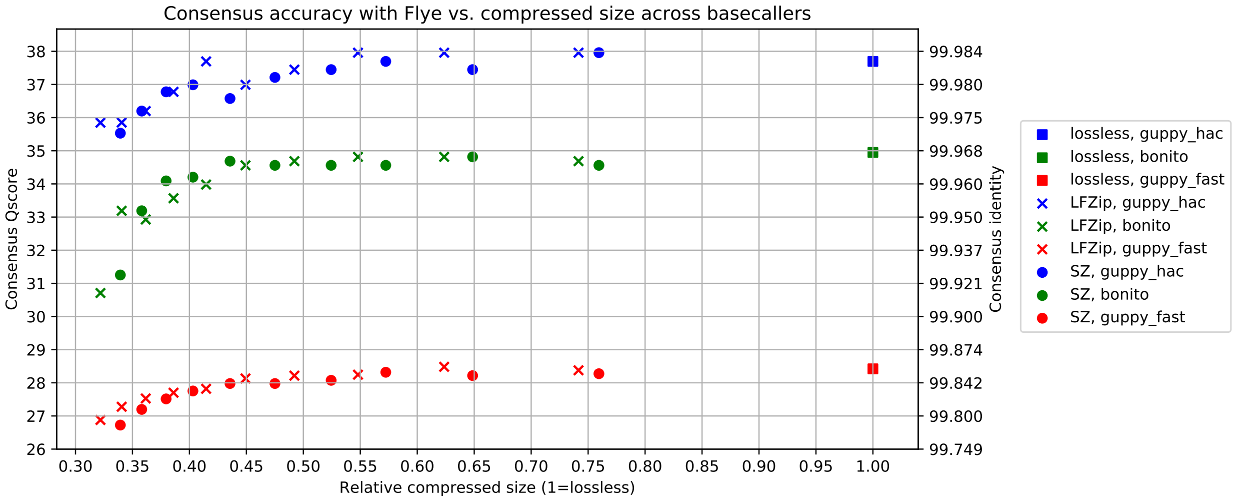

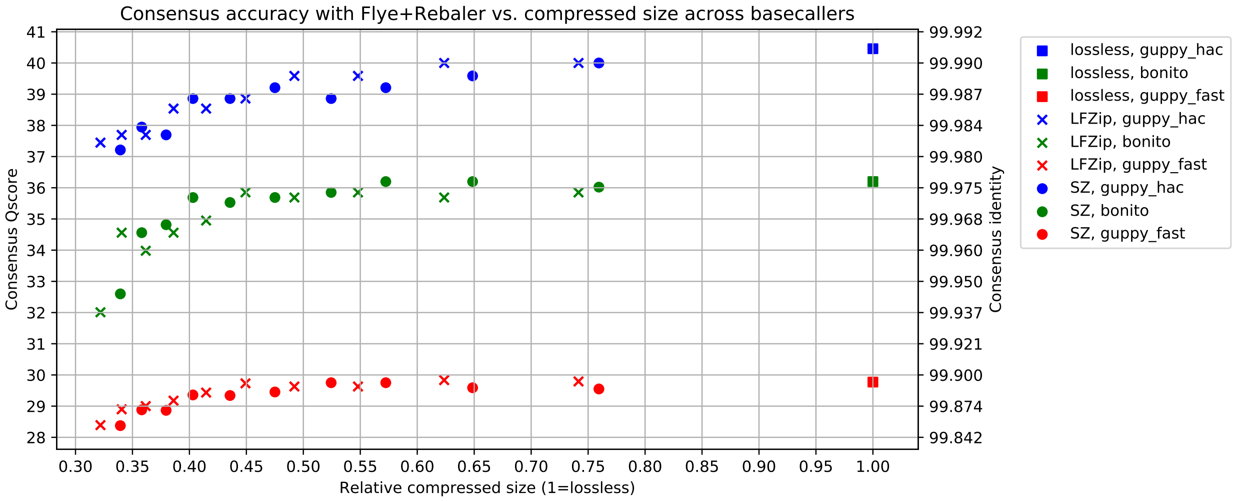

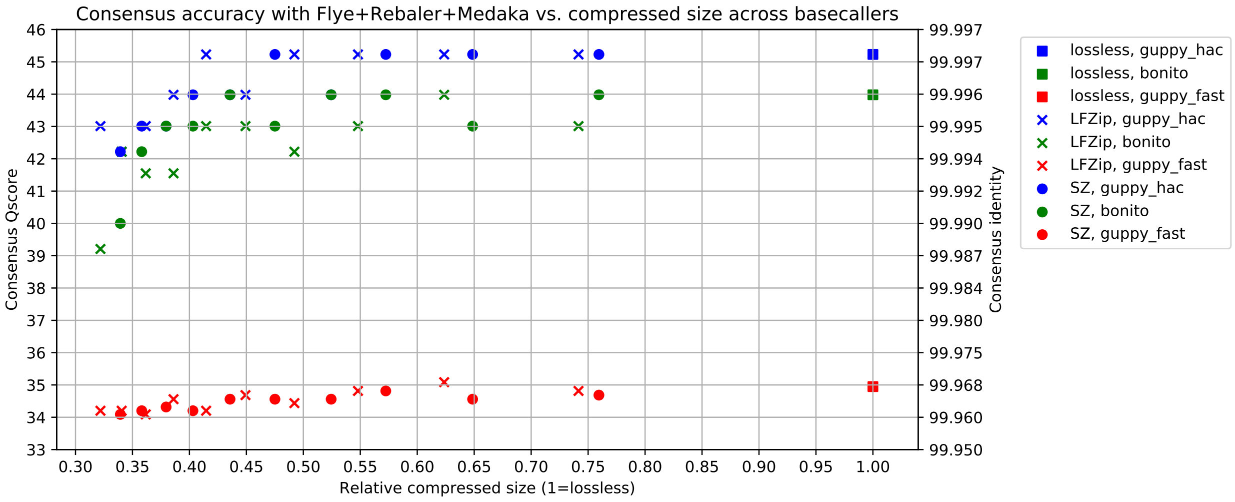

##### 2.1.4 Assembly accuracy across assembly stages

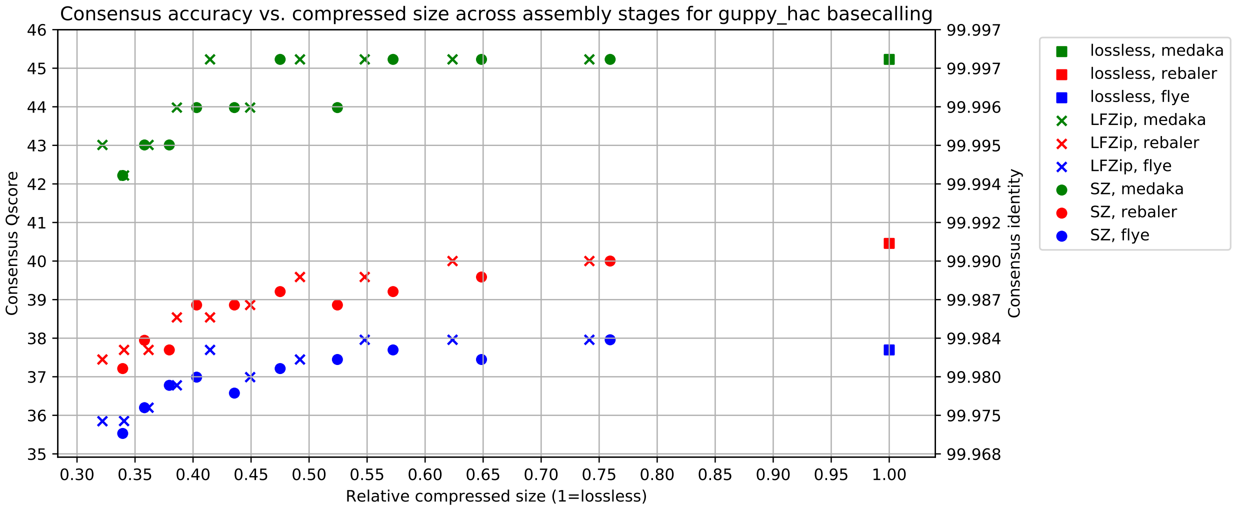

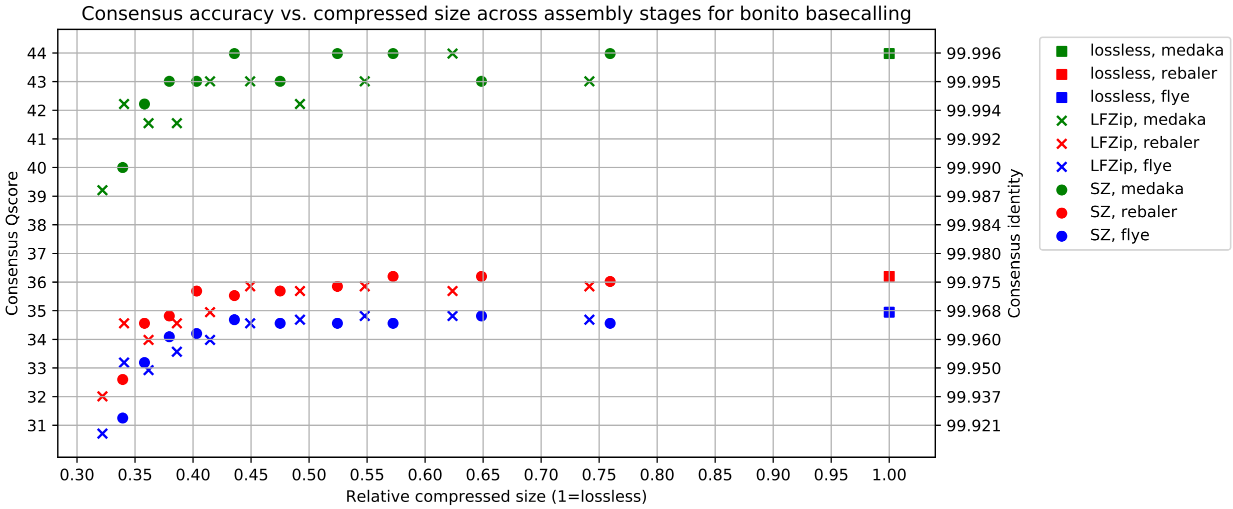

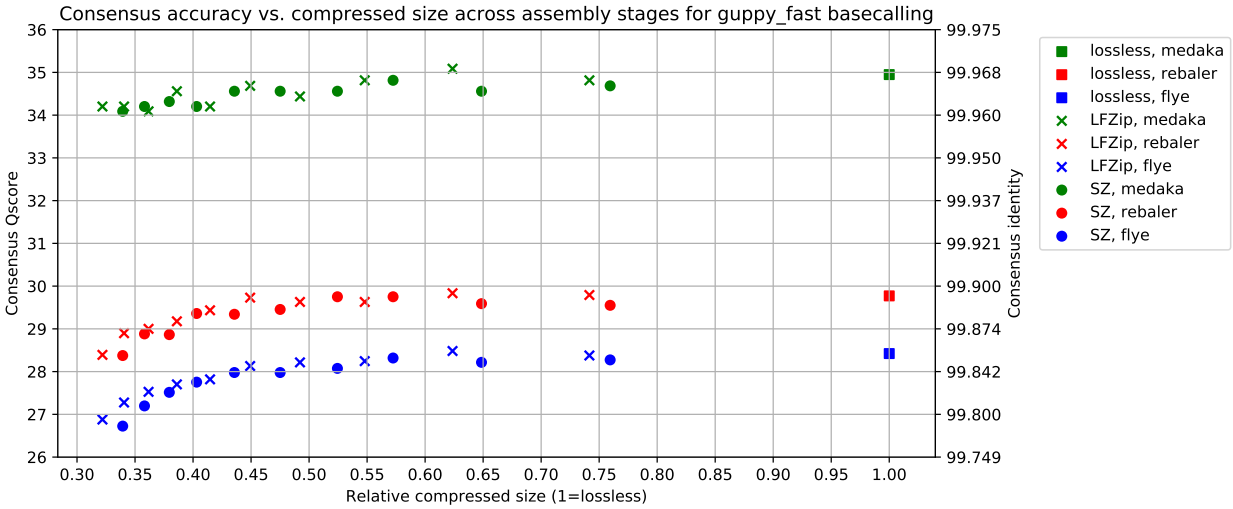

##### 2.1.5 Assembly accuracy across subsampling

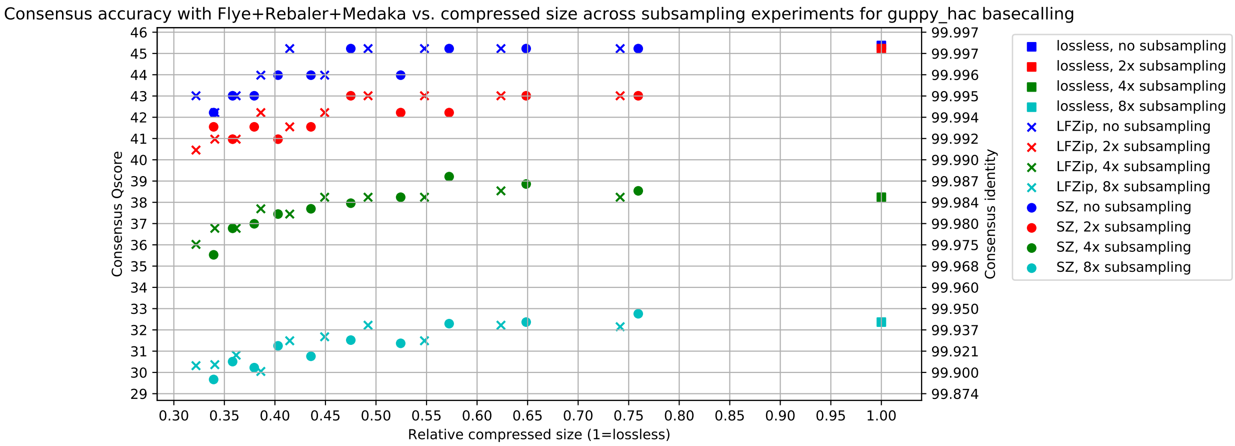

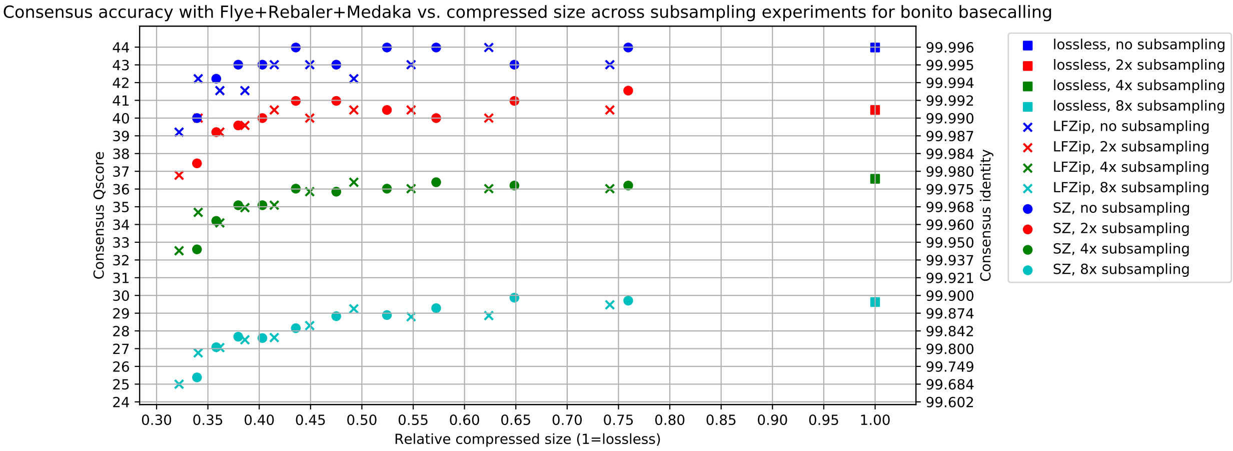

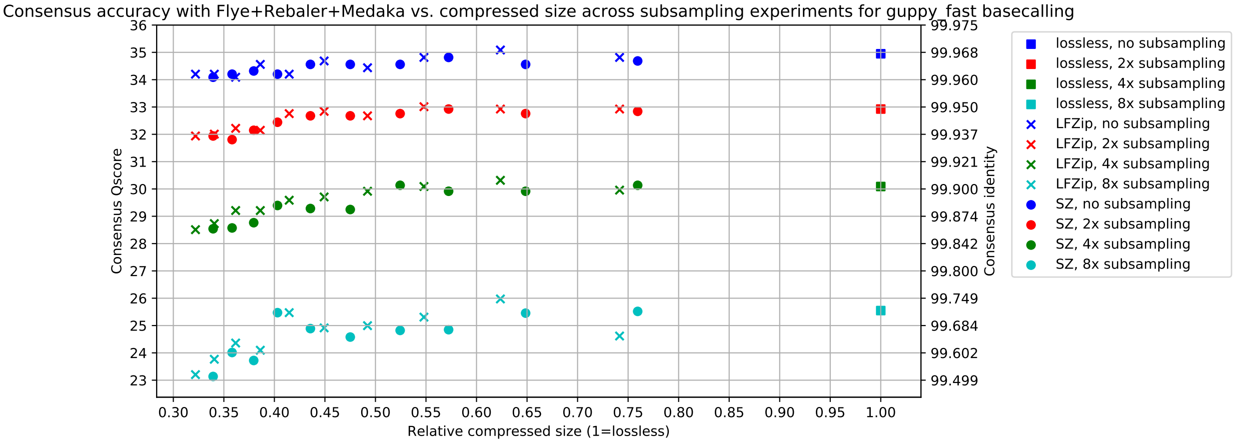

##### 2.1.6 Contig sizes and homopolymer accuracy

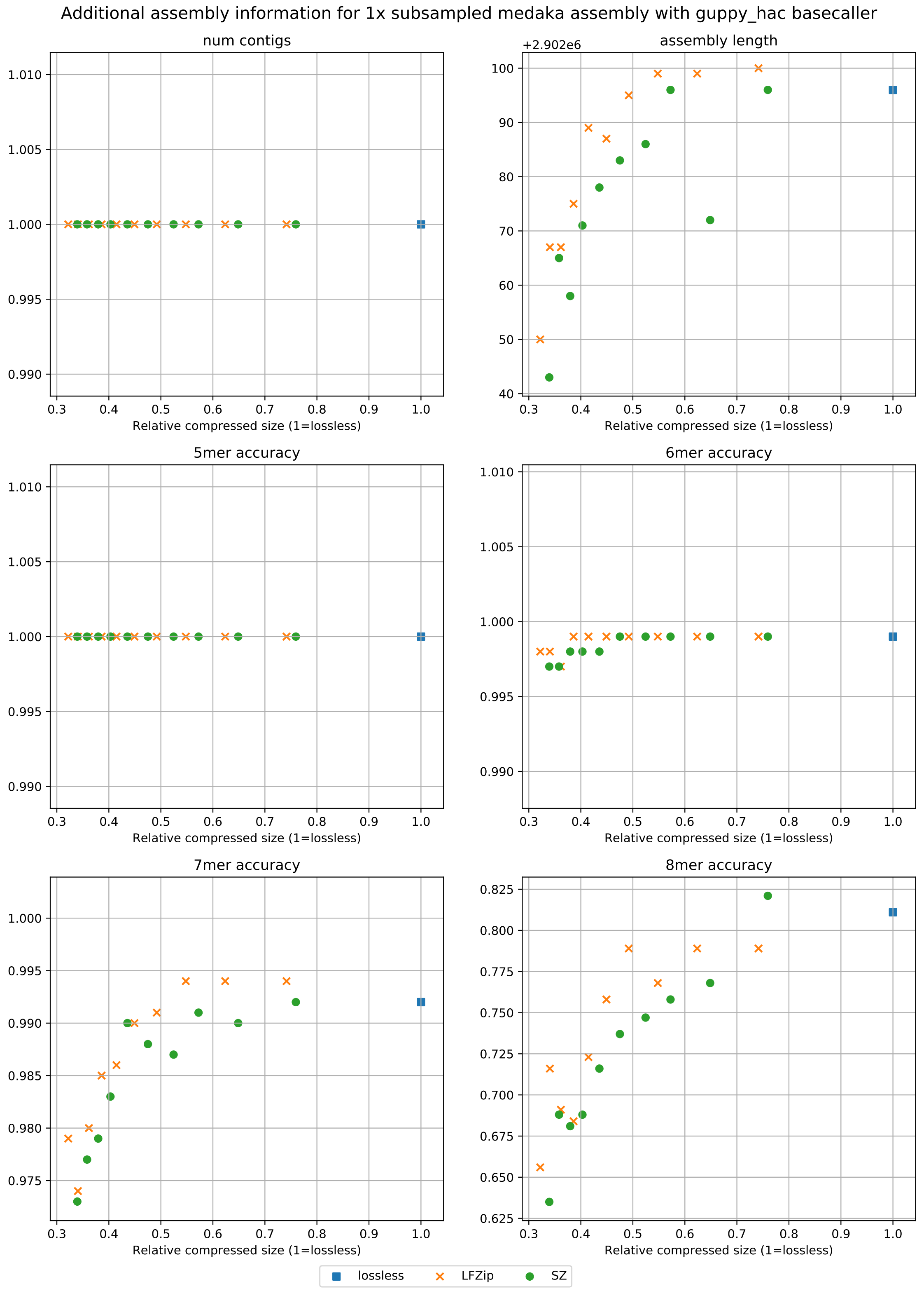

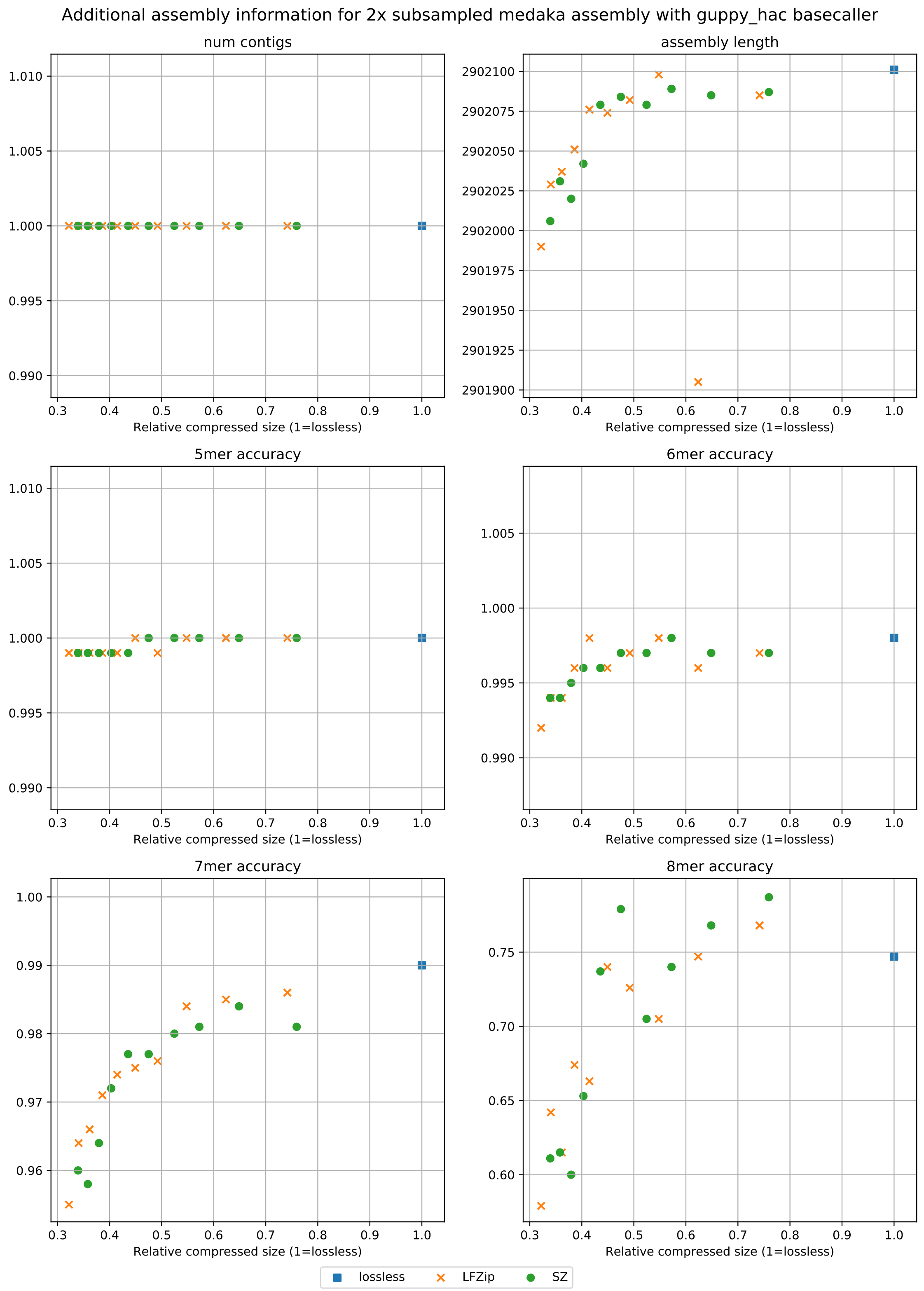
Additional assembly information for 2x subsampled medaka assembly with guppy_hac basecaller
num contigs
assembly length
5mer accuracy
6mer accuracy
7mer accuracy
8mer accuracy
Relative compressed size (1=lossless)
lossless
LFZip
SZ

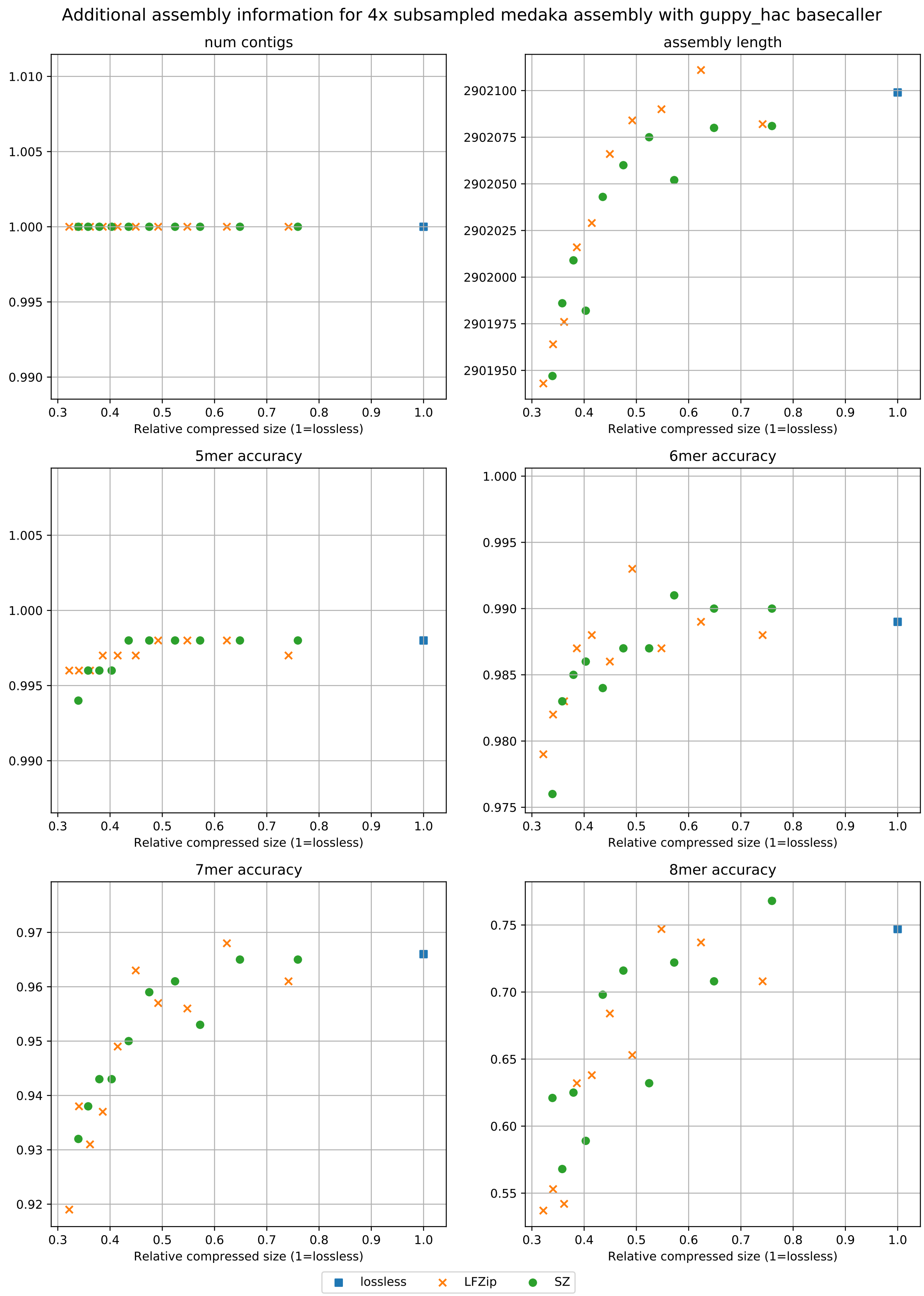

Additional assembly information for 4x subsampled medaka assembly with guppy_hac basecaller
num contigs
assembly length
5mer accuracy
6mer accuracy
7mer accuracy
8mer accuracy
Relative compressed size (1=lossless)
lossless
LFZip
SZ

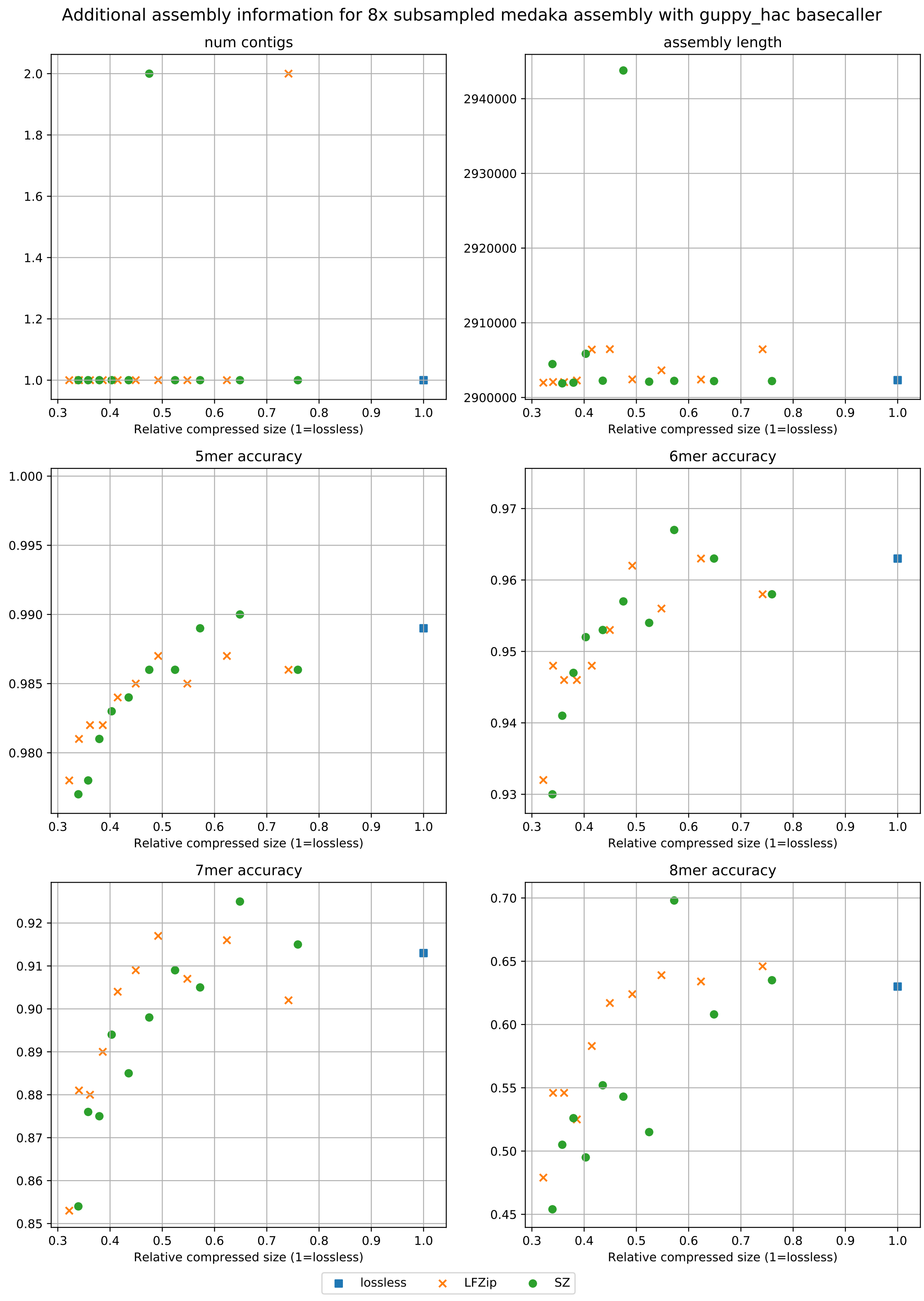

Additional assembly information for 8x subsampled medaka assembly with guppy_hac basecaller
num contigs
assembly length
5mer accuracy
6mer accuracy
7mer accuracy
8mer accuracy
Relative compressed size (1=lossless)
lossless
LFZip
SZ

#### 2.2 *K. pneumoniae*

##### 2.2.1 Lossy compression sizes

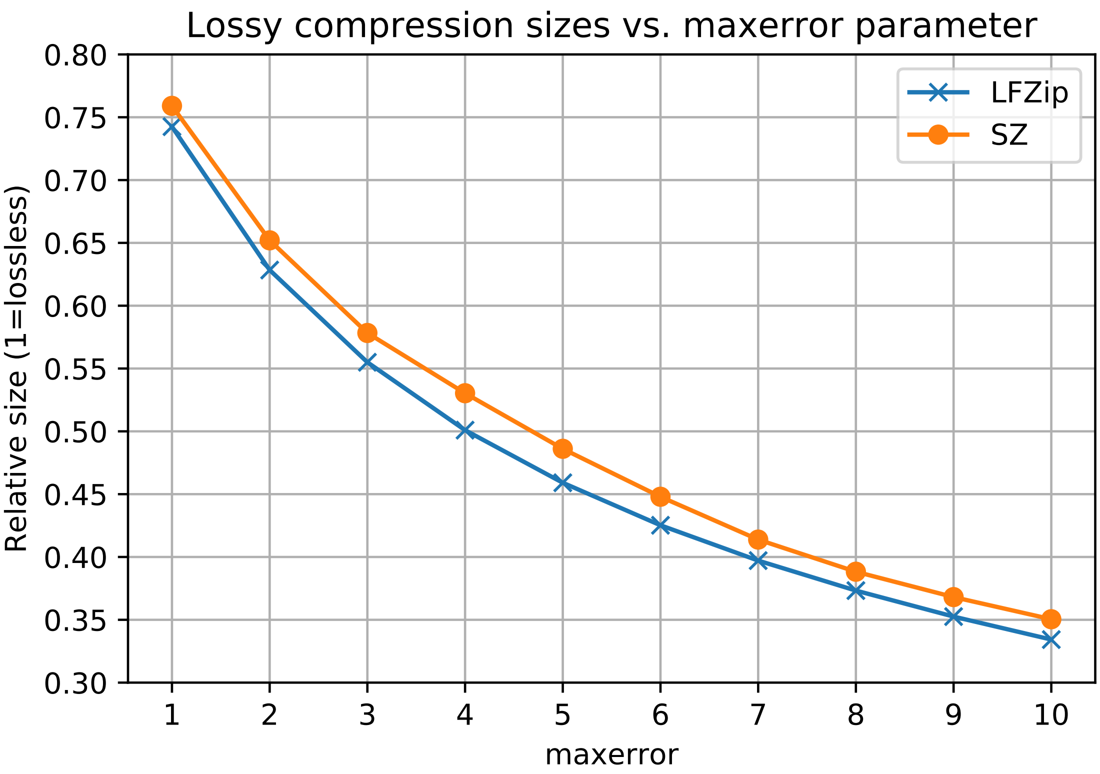

##### 2.2.2 Basecalling accuracy

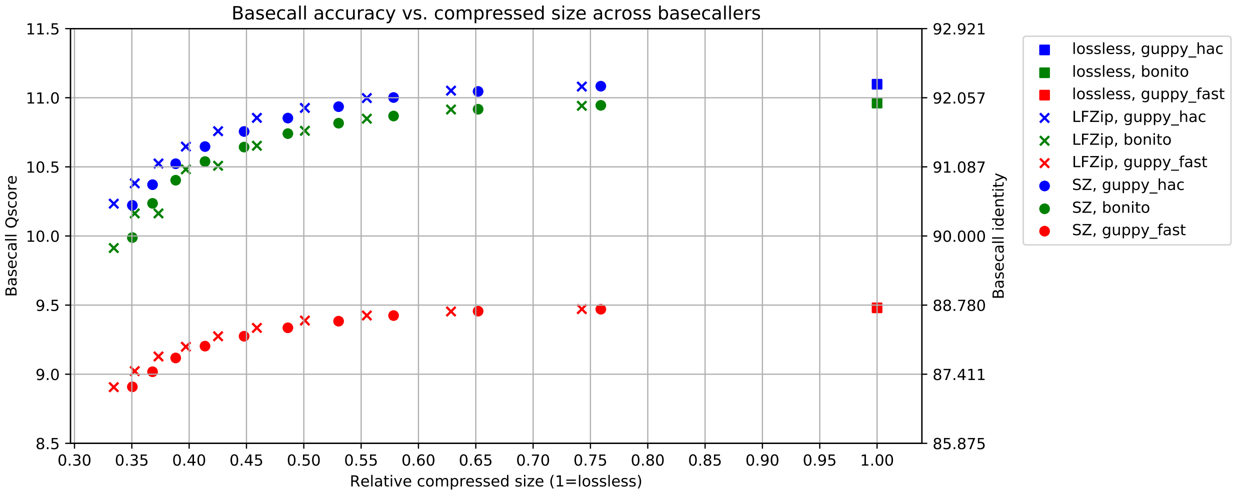

##### 2.2.3 Assembly accuracy across basecallers

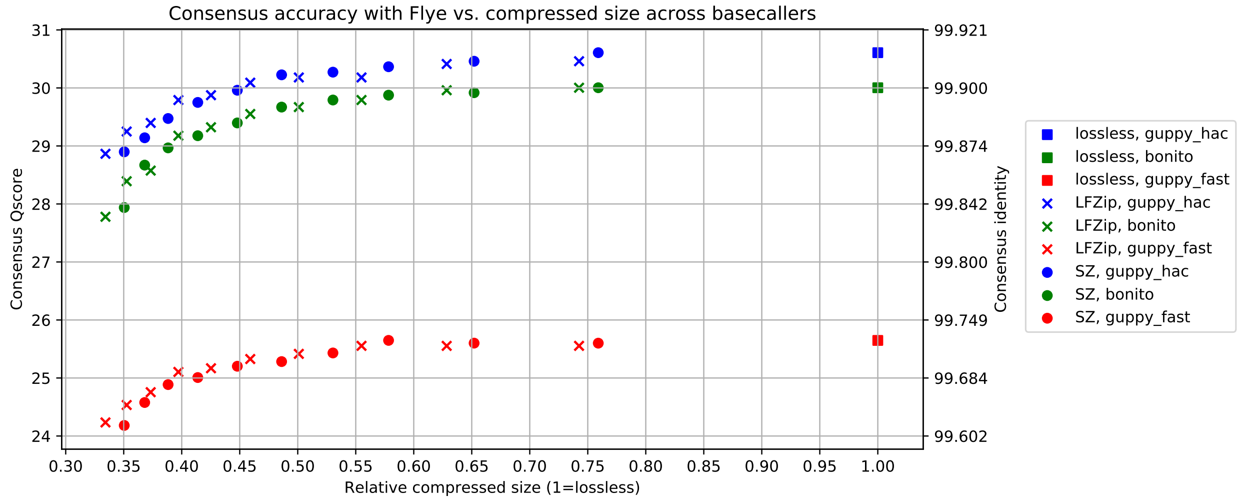

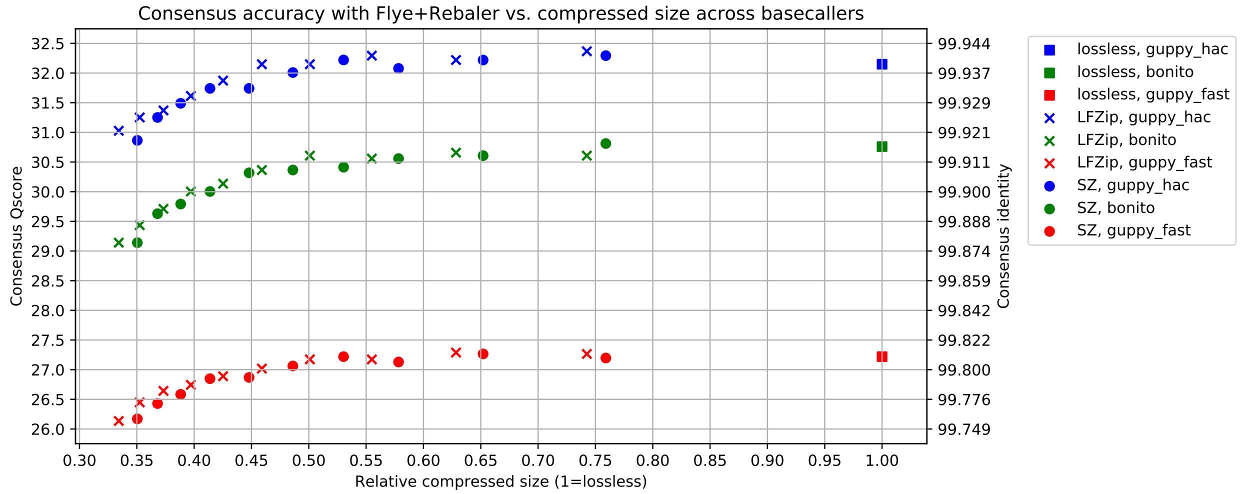

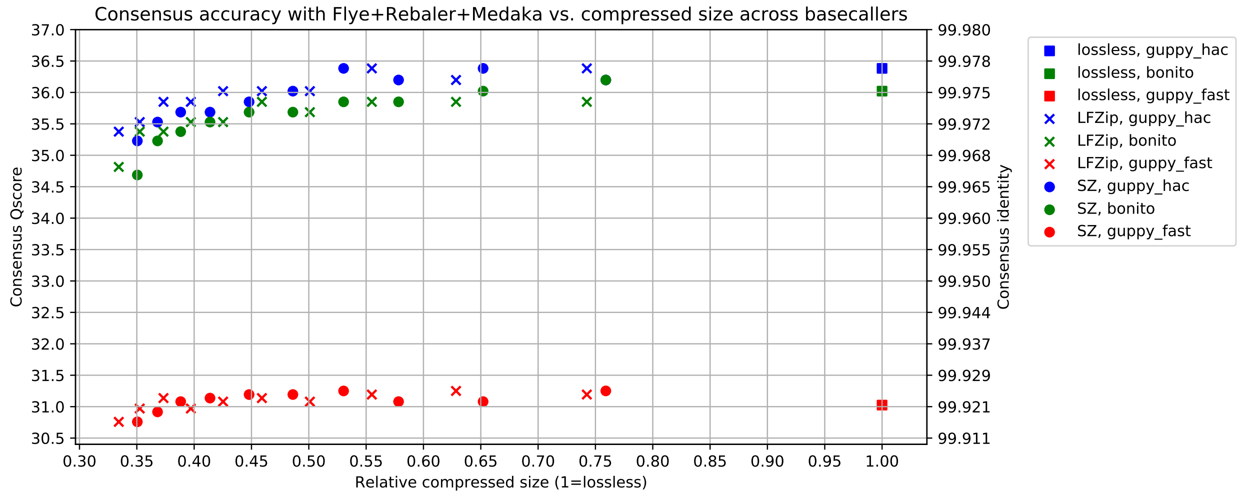

##### 2.2.4 Assembly accuracy across assembly stages

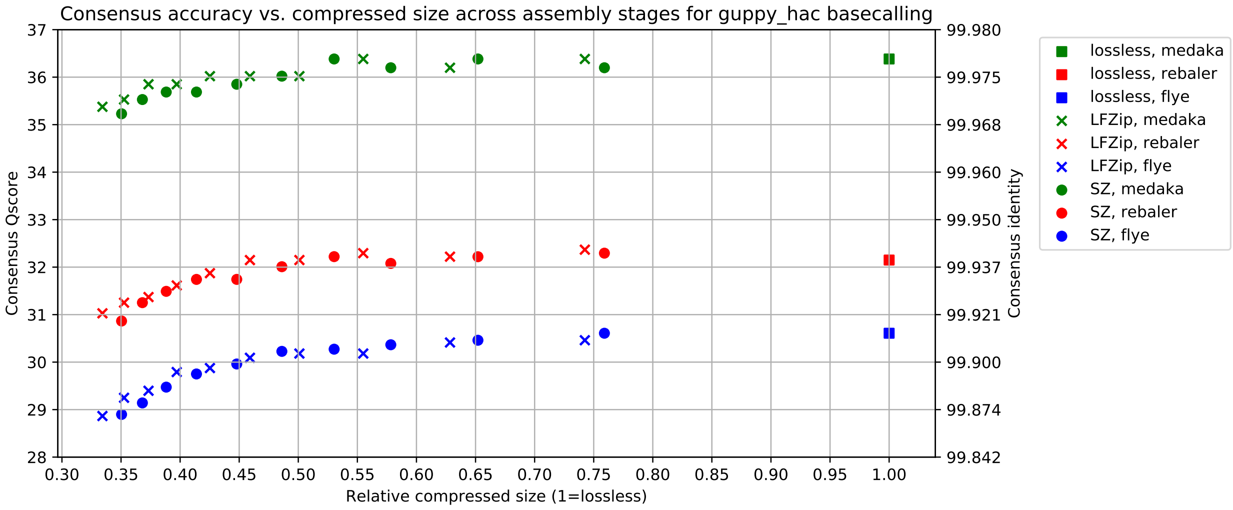

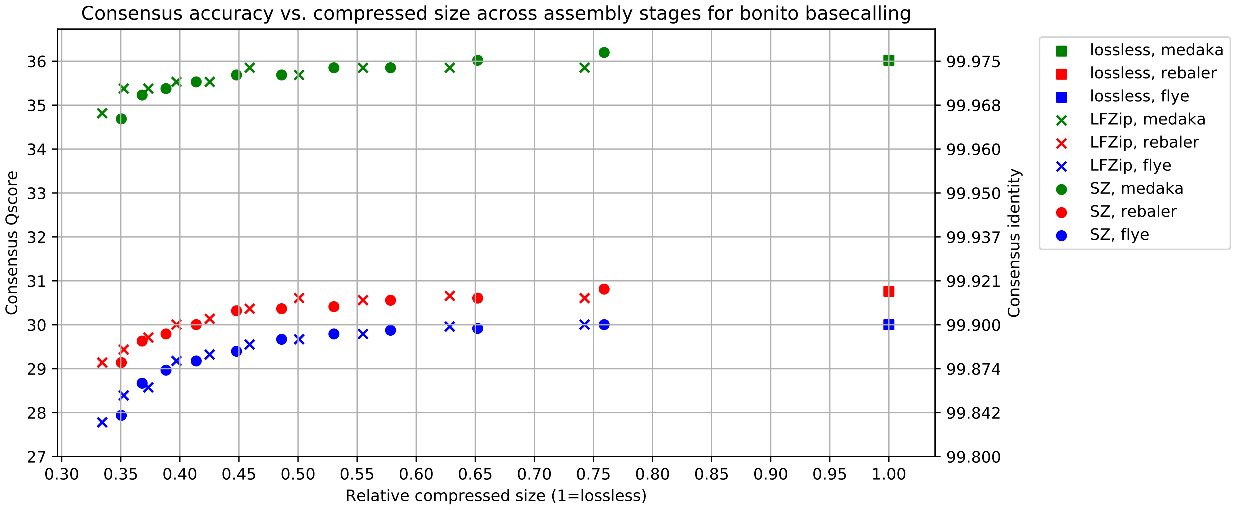

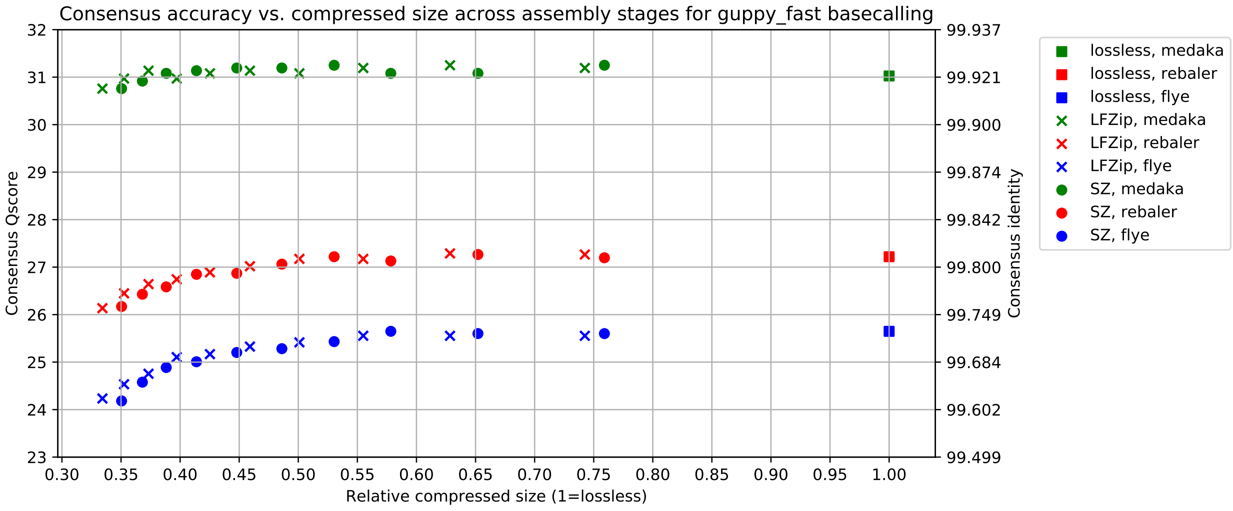

##### 2.2.5 Assembly accuracy across subsampling

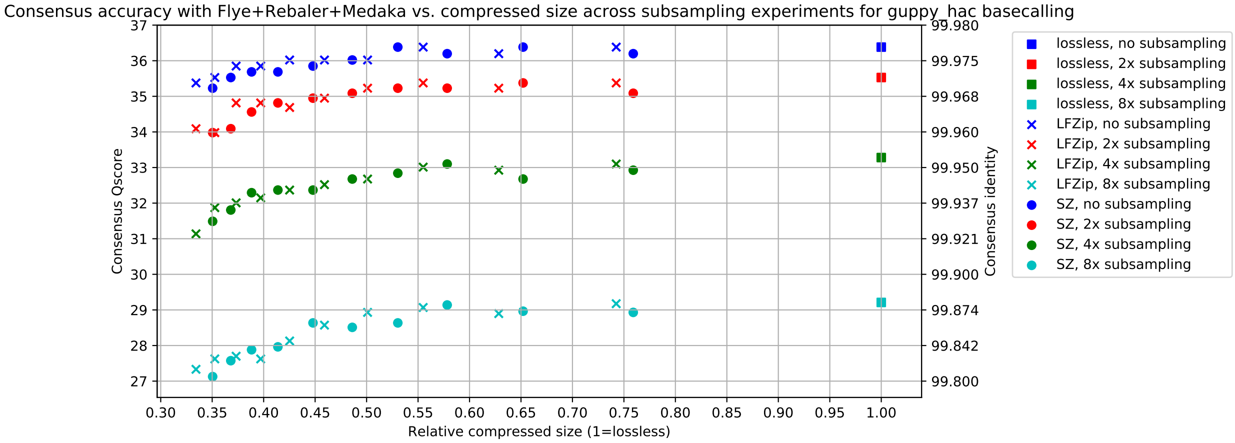

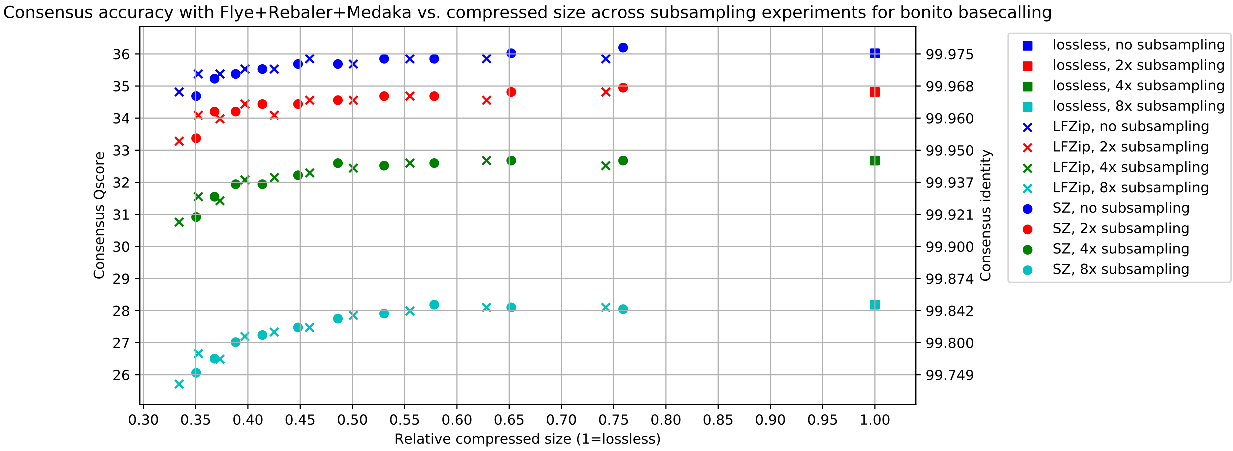

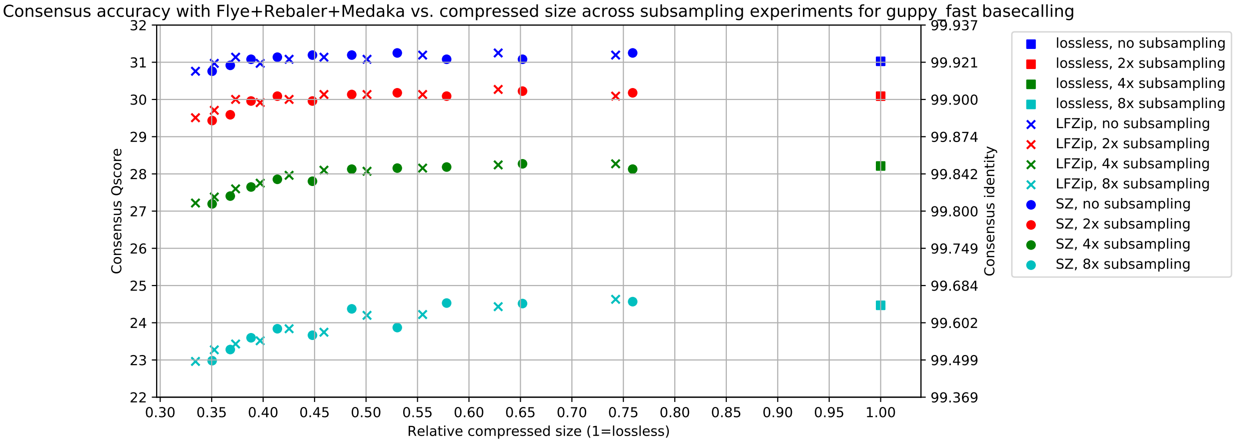

##### 2.2.6 Contig sizes and homopolymer accuracy

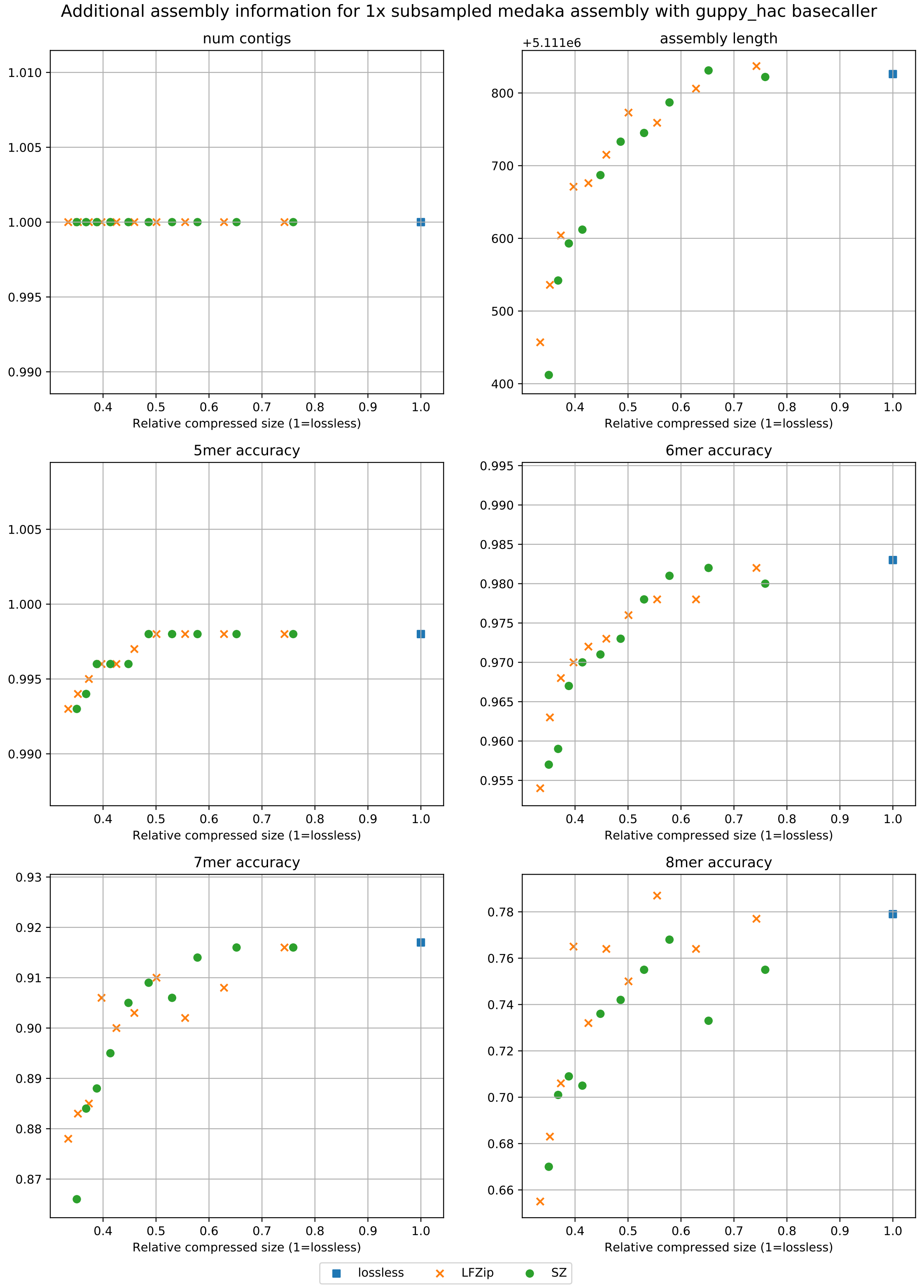

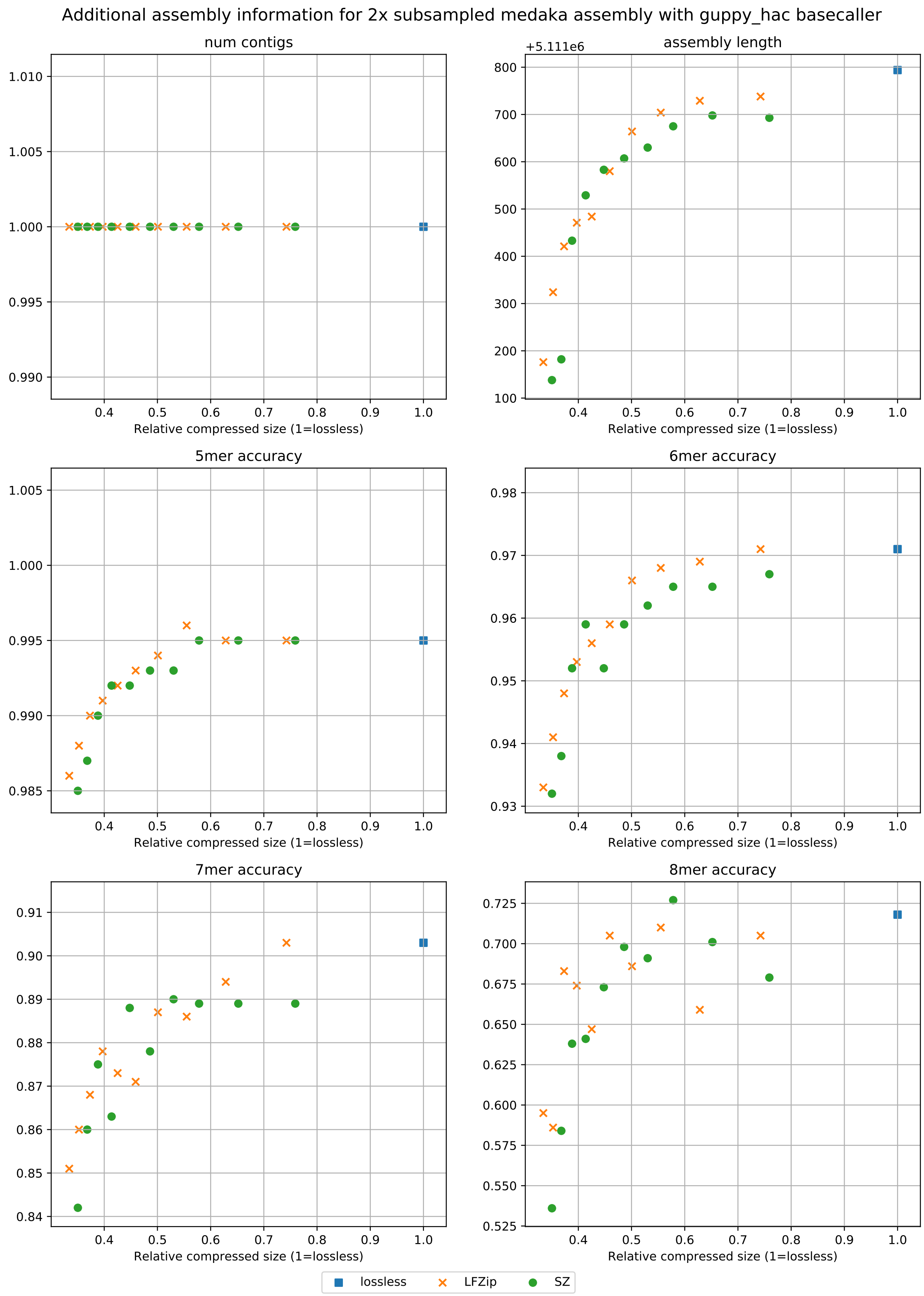

Additional assembly information for 2x subsampled medaka assembly with guppy_hac basecaller
num contigs
assembly length
+5.111e6
5mer accuracy
6mer accuracy
7mer accuracy
8mer accuracy
Relative compressed size (1=lossless)
lossless
LFZip
SZ

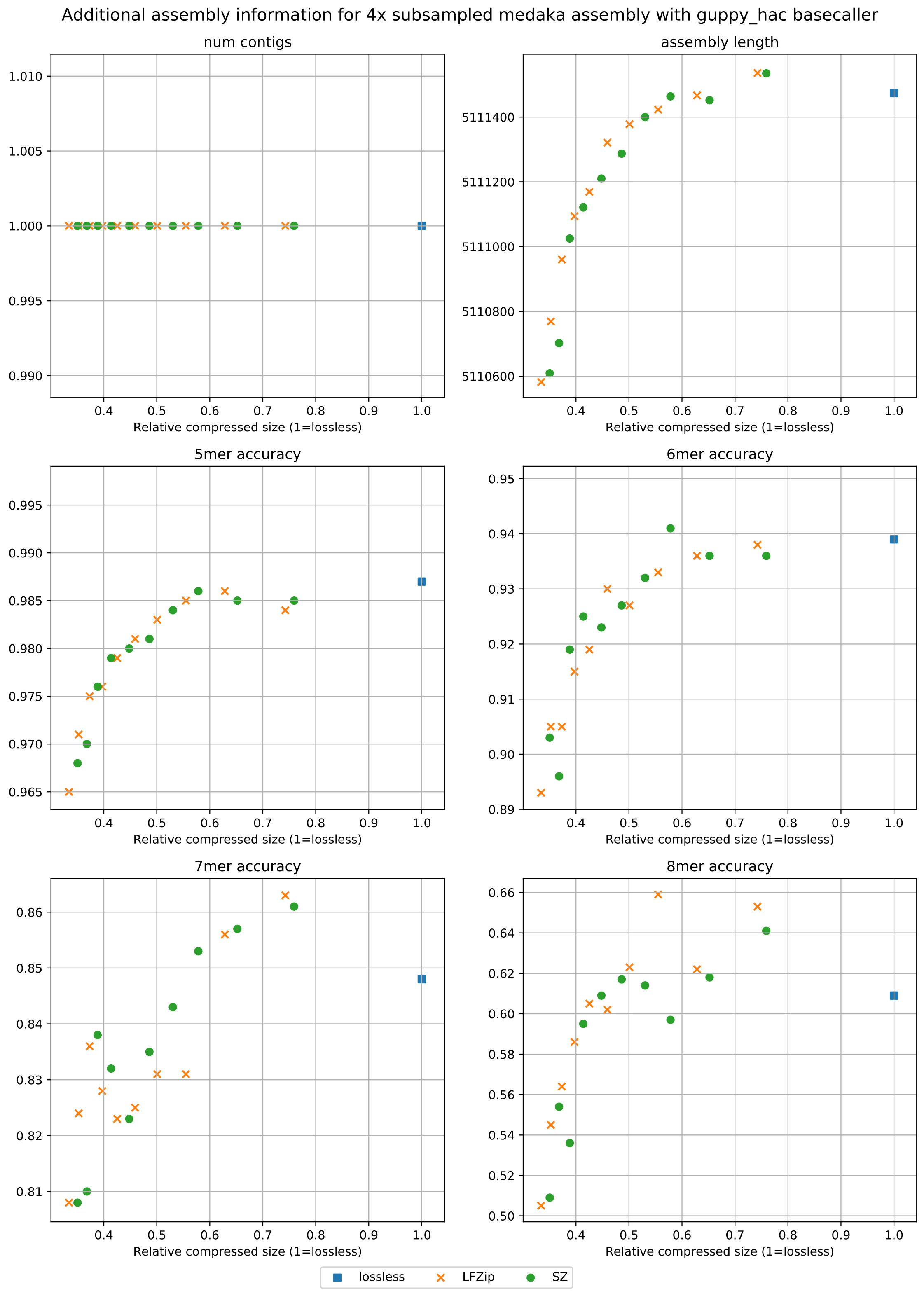

Additional assembly information for 4x subsampled medaka assembly with guppy_hac basecaller
num contigs
assembly length
5mer accuracy
6mer accuracy
7mer accuracy
8mer accuracy
Relative compressed size (1=lossless)
1.010
1.005
1.000
0.995
0.990
5111400
5111200
5111000
5110800
5110600
0.995
0.990
0.985
0.980
0.975
0.970
0.965
0.95
0.94
0.93
0.92
0.91
0.90
0.89
0.86
0.85
0.84
0.83
0.82
0.81
0.66
0.64
0.62
0.60
0.58
0.56
0.54
0.52
0.50
0.4
0.5
0.6
0.7
0.8
0.9
1.0
lossless
LFZip
SZ

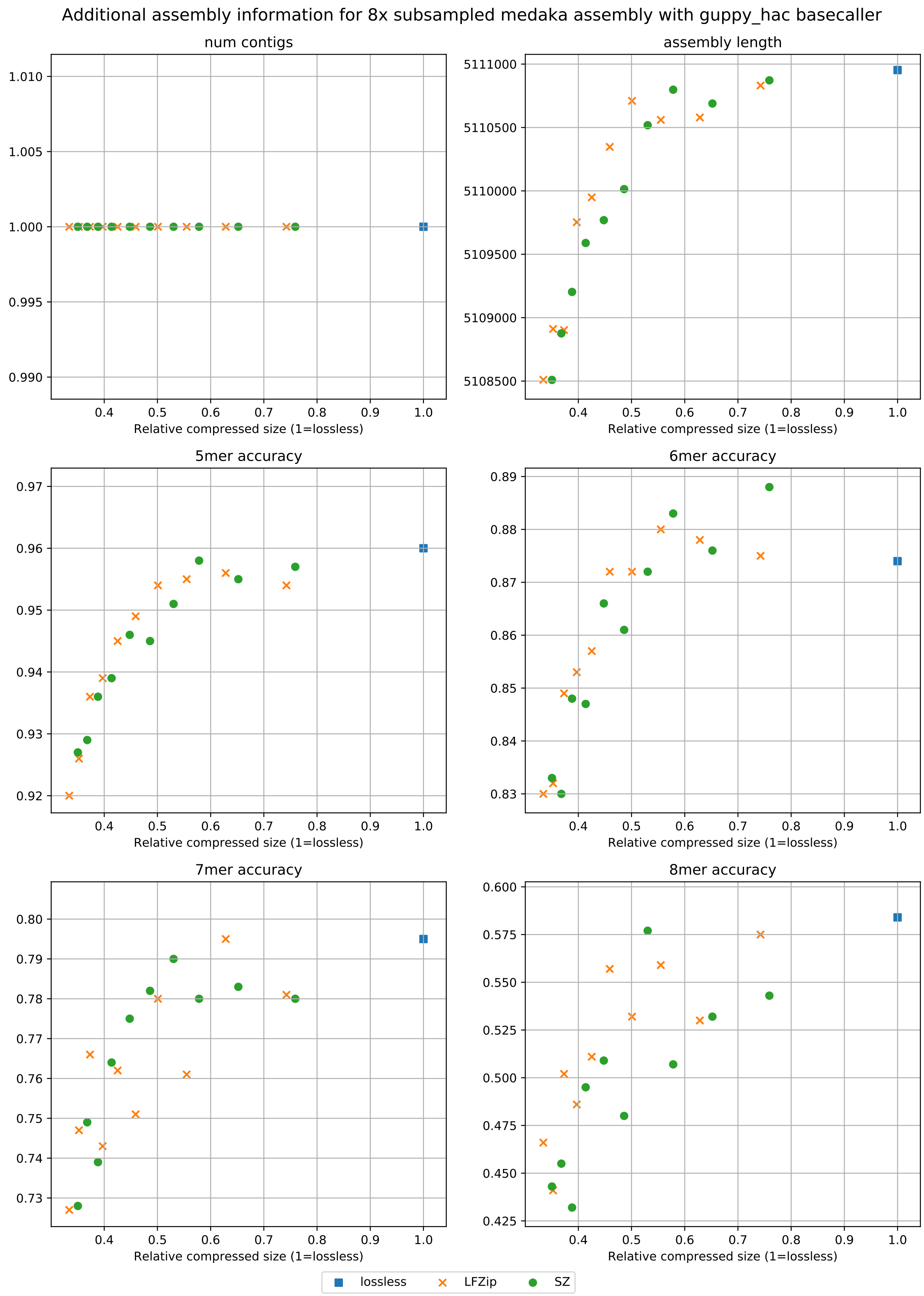

Additional assembly information for 8x subsampled medaka assembly with guppy_hac basecaller
num contigs
assembly length
5mer accuracy
6mer accuracy
7mer accuracy
8mer accuracy
Relative compressed size (1=lossless)
lossless
LFZip
SZ

##### 2.3 *E. coli*

###### 2.3.1 Lossy compression sizes

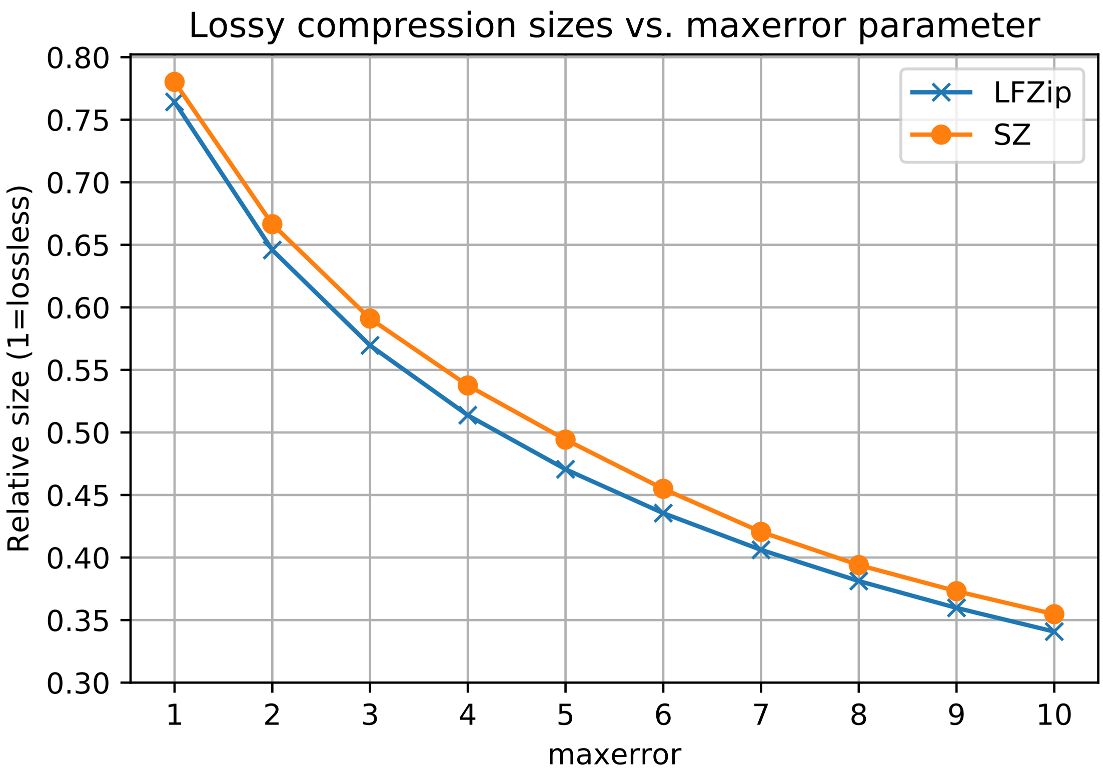

##### 2.3.2 Basecalling accuracy

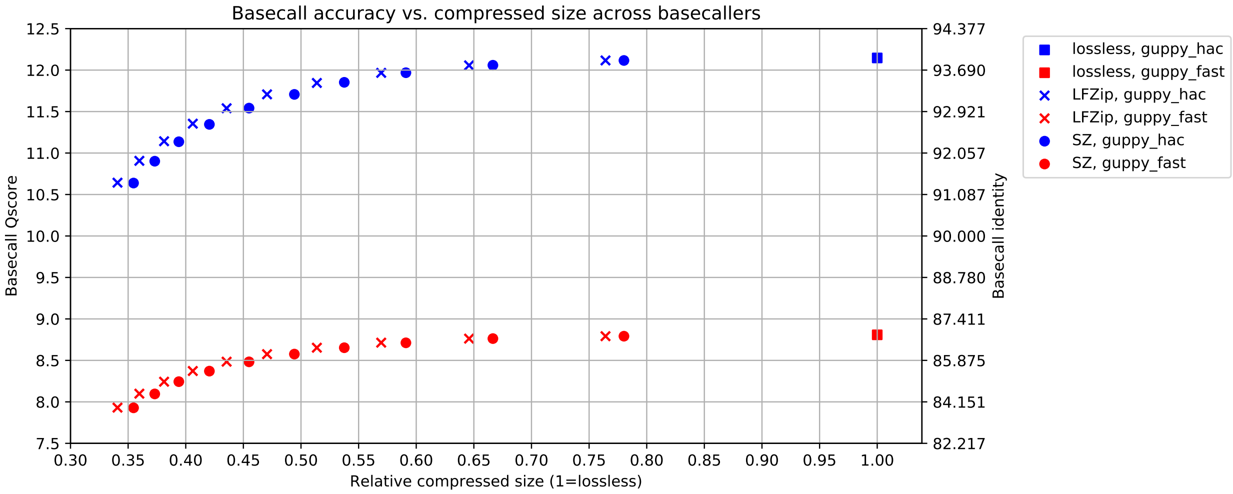

##### 2.3.3 Assembly accuracy across basecallers

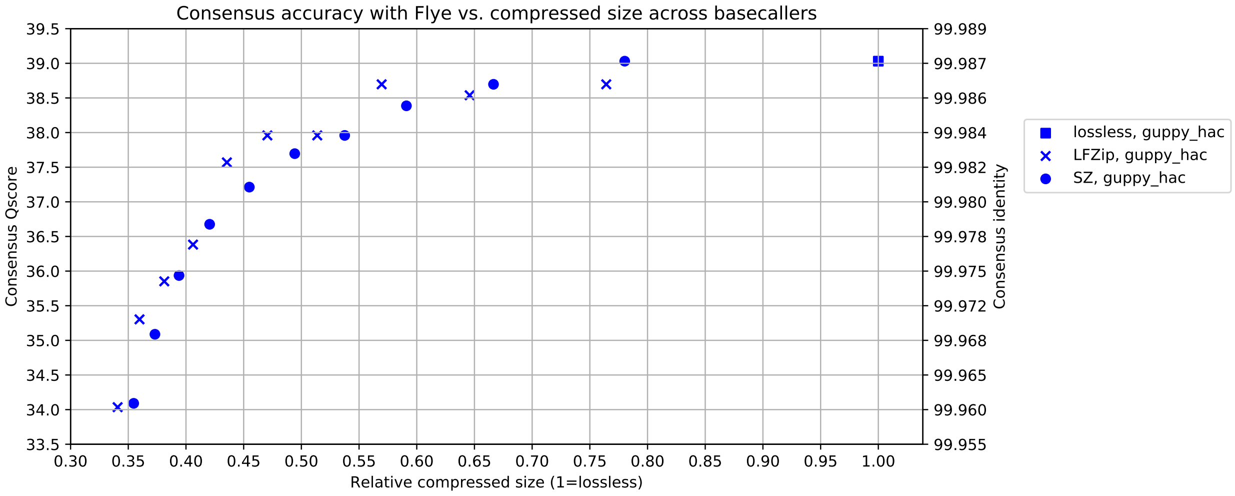

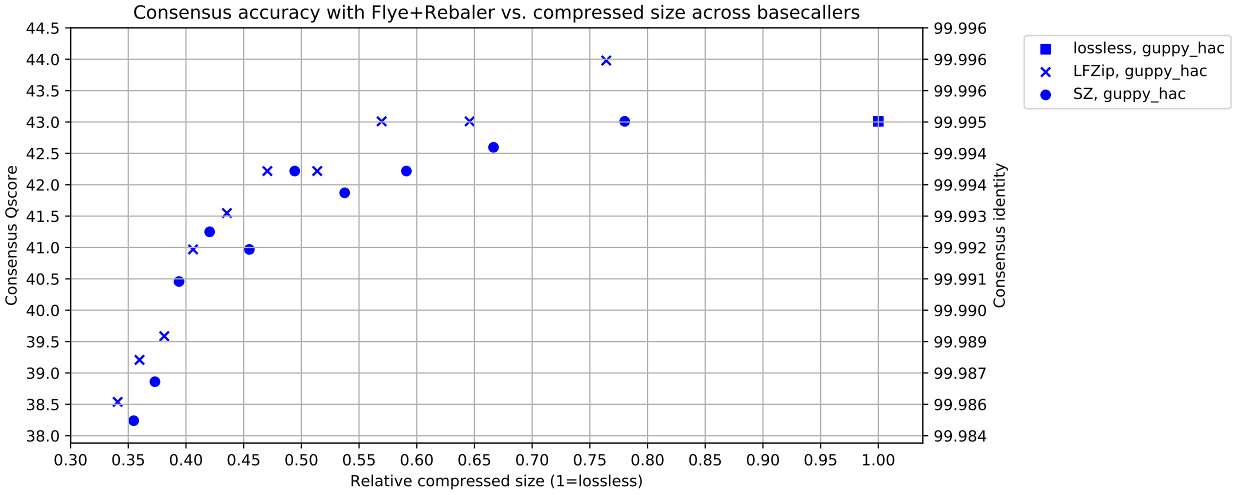

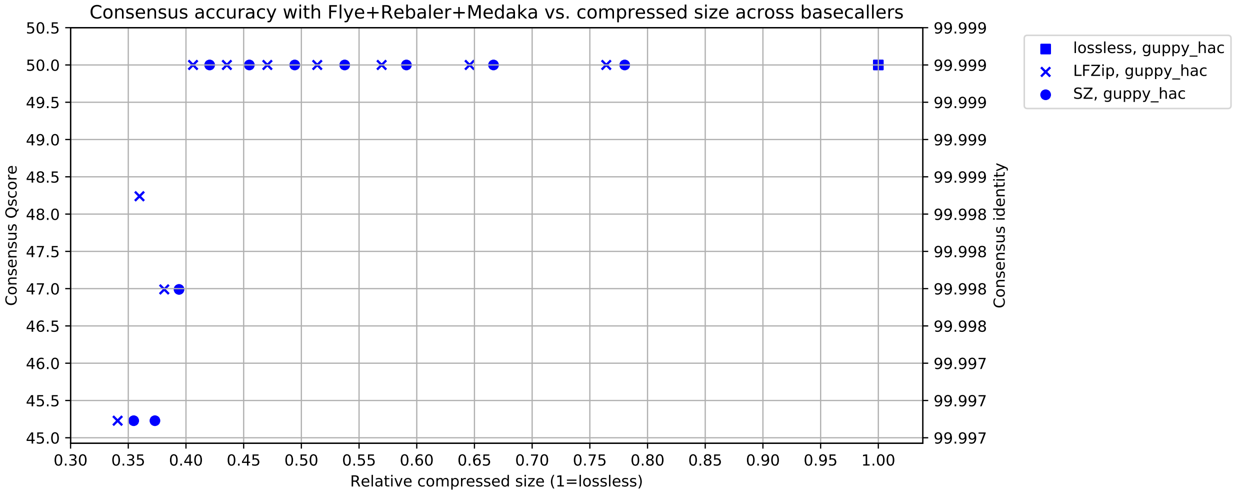

##### 2.3.4 Assembly accuracy across assembly stages

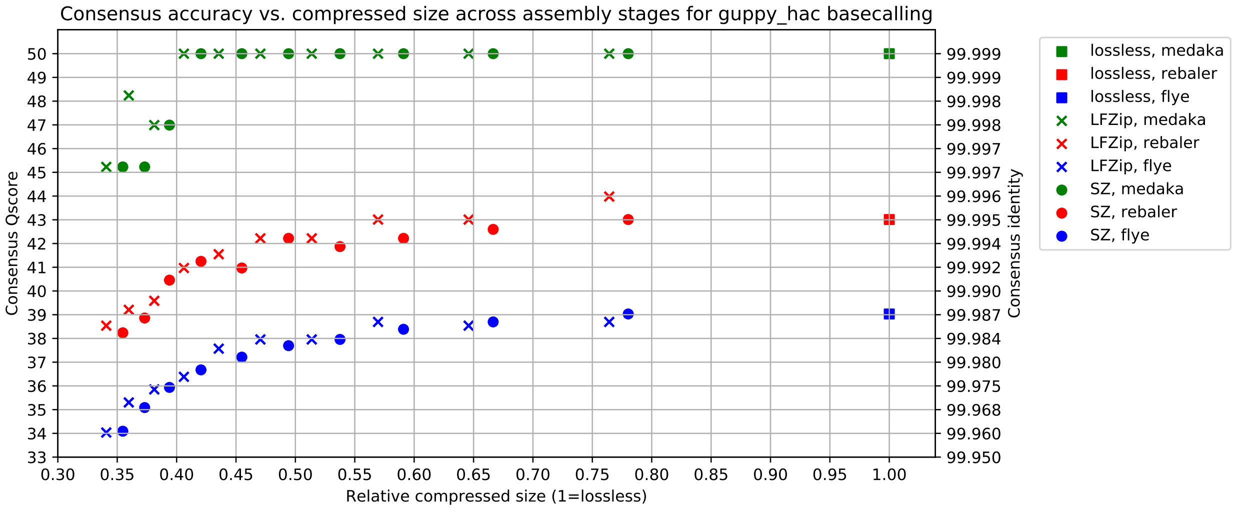

##### 2.3.5 Assembly accuracy across subsampling

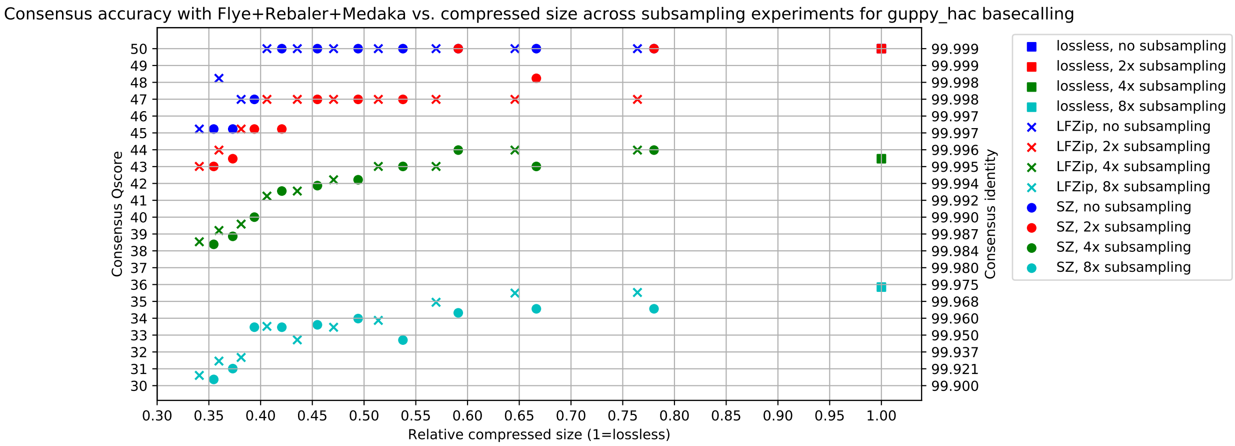

##### 2.3.6 Contig sizes and homopolymer accuracy

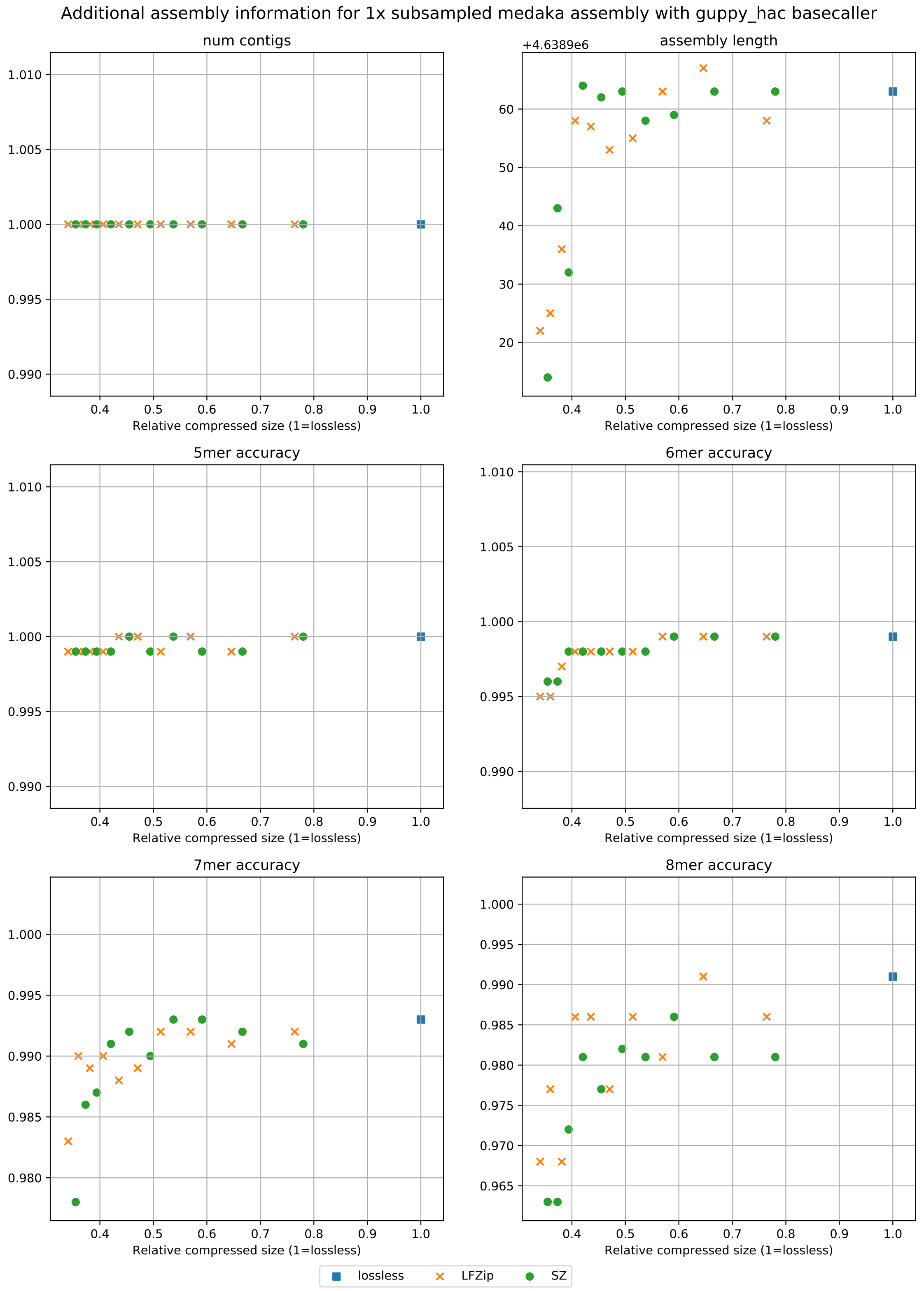

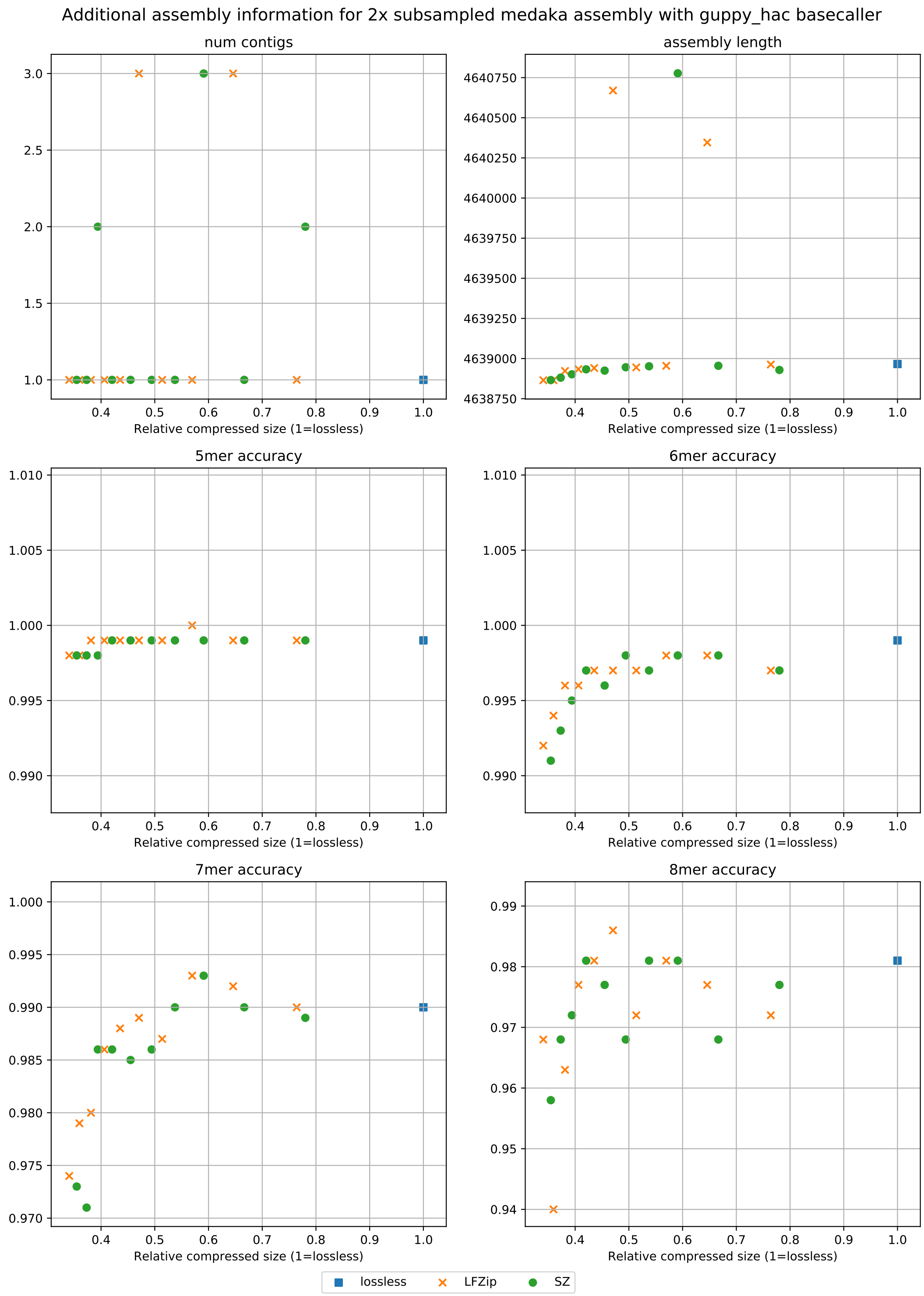

**Note on 2x subsampled assembly:** Although the num_contigs plot seems to show that lossy compression leads to fragmented assemblies, we found that $> 99.9\%$ of the total assembly length is contained in the largest contig for all compression parameters.

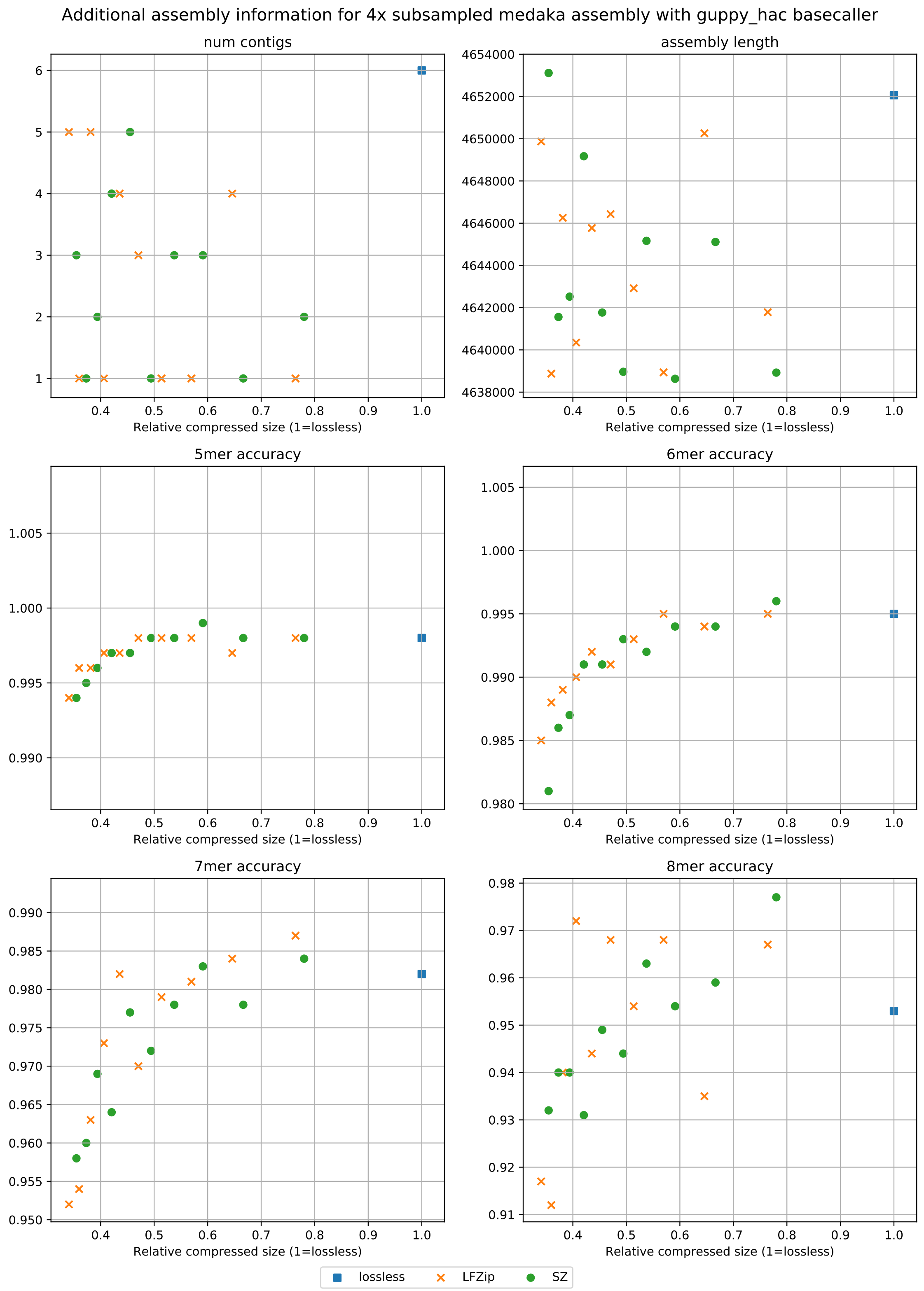
Additional assembly information for 4x subsampled medaka assembly with guppy_hac basecaller
num contigs
assembly length
5mer accuracy
6mer accuracy
7mer accuracy
8mer accuracy
Relative compressed size (1=lossless)
lossless
LFZip
SZ

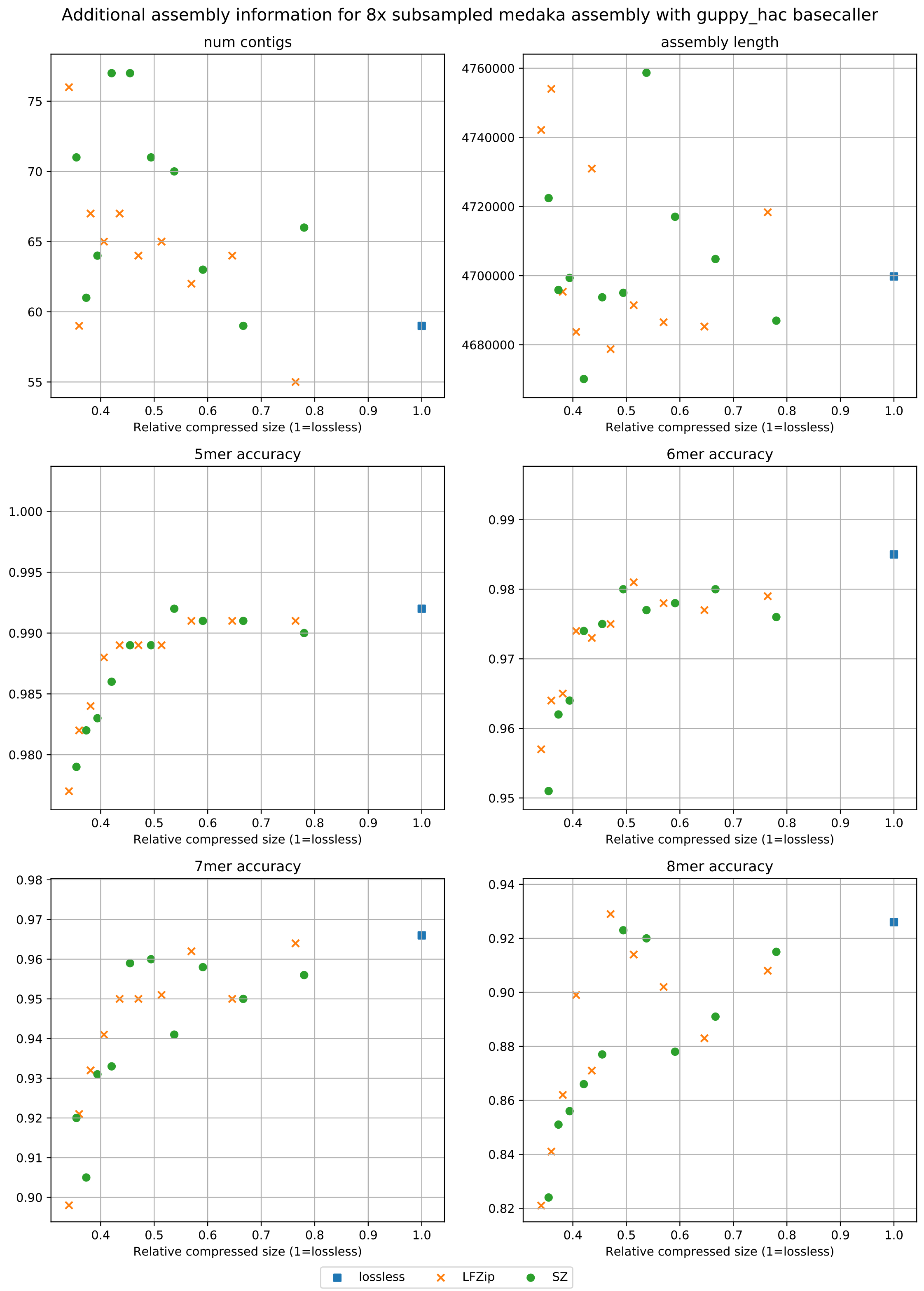

Additional assembly information for 8x subsampled medaka assembly with guppy_hac basecaller
num contigs
assembly length
5mer accuracy
6mer accuracy
7mer accuracy
8mer accuracy
Relative compressed size (1=lossless)
lossless
LFZip
SZ

#### 2.4 *H. sapiens*

##### 2.4.1 Lossy compression sizes

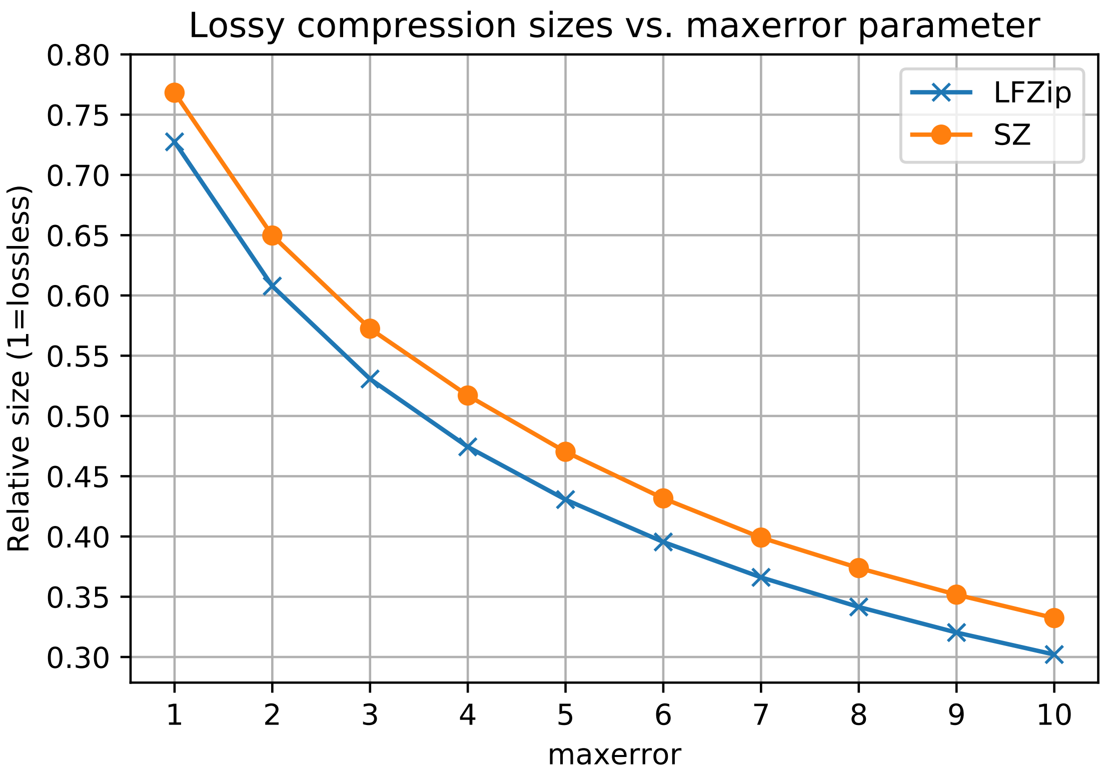

##### 2.4.2 Basecalling accuracy

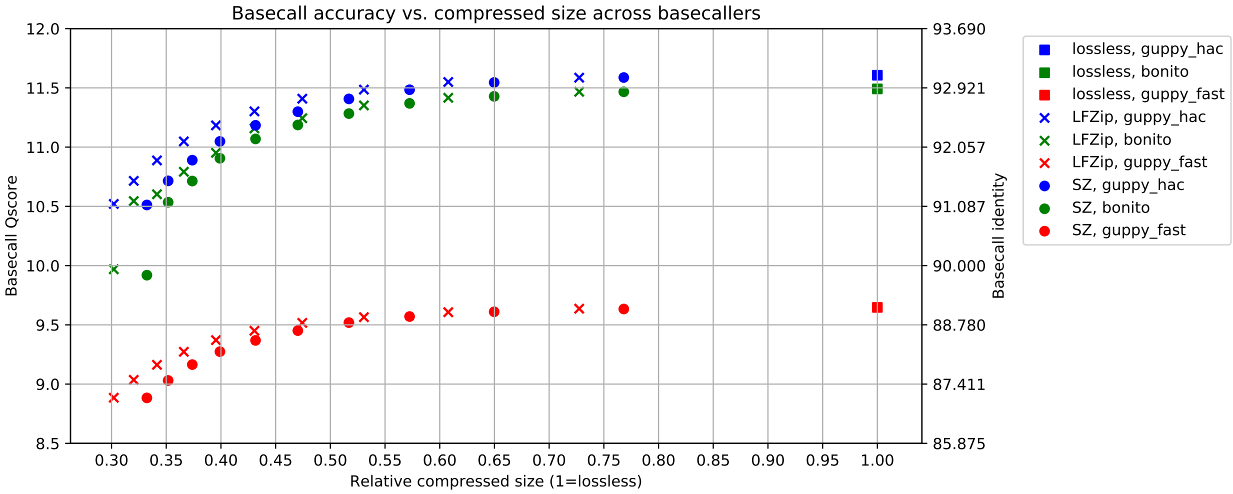

##### 2.4.3 Methylation accuracy

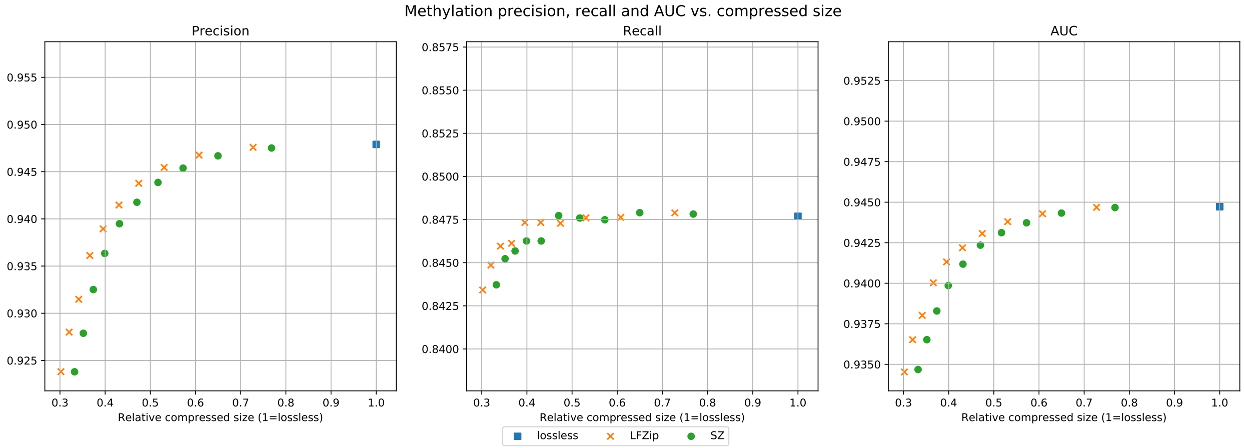

### 3 Instructions for installing tools

We list below are the steps for installing all the tools required for reproducing the results in the study. We will assume everything is installed in a working directory `$WORKINGDIR` and this variable is set using `export WORKINGDIR=/MY/PATH`. All experiments were performed on an Ubuntu 18.04.4 server with 40 Intel Xeon processors (2.2 GHz), 260 GB RAM and 8 Nvidia TITAN X (Pascal) GPUs. The default Python version was 3.7.6 (Anaconda).

```
cd $WORKINGDIR
```

**Clone git repository**

```
git clone https://github.com/shubhamchandak94/lossy_compression_evaluation
```

**Create and activate virtual environment used for running all the code**

```
python3 -m venv lossy_comp_env
source lossy_comp_env/bin/activate
pip install -q --upgrade pip
```

#### 3.1 Utility libraries

**ont_fast5_api**

Utility functions for conversion from multi to single read fast5.

```
pip install ont_fast5_api
```

**h5py**

For accessing fast5 files with raw data.

```
pip install h5py
```

**Seqtk**

Library used to manipulate basecalled fastq files.

```
wget https://github.com/lh3/seqtk/archive/v1.3.tar.gz
cd seqtk-1.3/
make
cd ../
rm v1.3.tar.gz
```

**Minimap2**

Aligner used by various tools in the pipeline.

```
wget https://github.com/lh3/minimap2/archive/v2.17.tar.gz
cd minimap2-2.17
make
cd ../
rm v2.17.tar.gz
```

##### Samtools

Tools for manipulating and analyzing SAM files containing aligned reads.

```
wget https://github.com/samtools/samtools/releases/download/1.10/samtools-1.10.tar.bz2
tar -xjvf samtools-1.10.tar.bz2
cd samtools-1.10/
./configure
make
cd ../
rm samtools-1.10.tar.bz2
```

##### fastmer

Library used for evaluating assemblies (specifically homopolymer accuracy).

```
pip install pyvcf
pip install pysam
git clone https://github.com/jts/assembly_accuracy
cd assembly_accuracy/
git checkout ff822506aa12958a203c093257cdbfcf7abd6308
cd ../
```

##### BCFtools

Used for generating fasta for NA12878 from reference fasta and GIAB VCF file.

```
wget https://github.com/samtools/bcftools/releases/download/1.11/bcftools-1.11.tar.bz2
tar -xjvf bcftools-1.11.tar.bz2
cd bcftools-1.11/
./configure
make
cd ../
rm bcftools-1.11.tar.bz2
```

#### 3.2 Compressors

##### LFZip

Lossy compressor.

```
conda create -n python3_6_env python=3.6
conda activate python3_6_env
conda config --add channels conda-forge
conda install lfzip=1.1
pip install h5py
conda deactivate
```

### SZ

Lossy compressor.

```
wget https://github.com/disheng222/SZ/archive/v2.1.8.3.tar.gz
tar -xzvf v2.1.8.3.tar.gz
cd SZ-2.1.8.3/
./configure --prefix=$WORKINGDIR/SZ-2.1.8.3
make
make install
```

```
cd ../
rm v2.1.8.3.tar.gz
```

###### VBZ python plugin

Lossless compressor.

```
conda activate python3_6_env # created above for lfzip
wget https://github.com/nanoporetech/vbz_compression/releases/download/v1.0.1/\
pyvbz-1.0.1-cp36-cp36m-linux_x86_64.whl
pip install pyvbz-1.0.1-cp36-cp36m-linux_x86_64.whl
conda deactivate
rm pyvbz-1.0.1-cp36-cp36m-linux_x86_64.whl
```

#### 3.3 Basecallers

###### Guppy

```
wget https://mirror.oxfordnanoportal.com/software/analysis/ont-guppy_3.6.1_linux64.tar.gz
tar -xzvf ont-guppy_3.6.1_linux64.tar.gz
rm ont-guppy_3.6.1_linux64.tar.gz
```

###### Bonito

```
pip install bonito==0.2.0
```

Medaka model for bonito:

```
mkdir bonito_medaka && cd bonito_medaka/
wget https://nanoporetech.box.com/shared/static/oukeesfjc6406t5po0x2hlw97lnelkyl.hdf5
cd ../
```

#### 3.4 Assembly/consensus

###### Flye

Assembler.

```
wget https://github.com/fenderglass/Flye/archive/2.7.1.tar.gz
tar -xzvf 2.7.1.tar.gz
cd Flye-2.7.1/
make
cd ../
rm 2.7.1.tar.gz
```

###### Racon

Consensus/polishing tool.

```
wget https://github.com/lbcb-sci/racon/releases/download/1.4.13/racon-v1.4.13.tar.gz
tar -xzvf racon-v1.4.13.tar.gz
cd racon-v1.4.13/
mkdir build/
cd build/
cmake -DCMAKE_BUILD_TYPE=Release ..
make
cd ../../
rm racon-v1.4.13.tar.gz
```

##### Rebaler

Performs several runs of Racon to polish assembly.

```
wget https://github.com/rrwick/Rebaler/archive/v0.2.0.tar.gz
tar -xzvf v0.2.0.tar.gz
rm v0.2.0.tar.gz
pip install biopython
```

##### Medaka

Performs further polishing of the Rebaler consensus using a neural network based approach.

```
git clone https://github.com/nanoporetech/medaka.git
cd medaka/
git checkout v1.0.3 # used v0.11.5 for guppy_fast due to errors in v1.0.3
conda activate python3_6_env  # created above for lfzip
make install
cd ../
conda deactivate
```

#### 3.5 Methylation calling

##### Megalodon

```
pip install megalodon==2.1.0
```

##### Rerio

Basecalling model for calling methylation at CpG motifs.

```
git clone https://github.com/nanoporetech/rerio
cd rerio
git checkout 82d5b18
./download_model.py basecall_models/res_dna_r941_min_modbases_5mC_CpG_v001
cd ../
```

### 4 Instructions for downloading datasets

The instructions for downloading the datasets used in the study are listed below. We will assume everything is stored in a working directory `$WORKINGDIR` and this variable is set using export `WORKINGDIR=/MY/PATH`.

```
cd $WORKINGDIR/
mkdir -p data/
cd data/
```

#### 4.1 *S. aureus*

```
mkdir Staphylococcus_aureus_CAS38_02/
cd Staphylococcus_aureus_CAS38_02/
mkdir fast5/ && cd fast5/
wget -O Staphylococcus_aureus_CAS38_02_fast5s.tar.gz \
https://bridges.monash.edu/ndownloader/files/14260568
tar -xzvf Staphylococcus_aureus_CAS38_02_fast5s.tar.gz
cd ../
wget -O Staphylococcus_aureus_CAS38_02_reference.fasta.gz \
https://bridges.monash.edu/ndownloader/files/14260241
gunzip Staphylococcus_aureus_CAS38_02_reference.fasta.gz
```

Cleanup:

```
rm fast5/Staphylococcus_aureus_CAS38_02_fast5s.tar.gz
mkdir experiments/
cd ../
```

#### 4.2 *K. pneumoniae*

```
mkdir Klebsiella_pneumoniae_INF032/
cd Klebsiella_pneumoniae_INF032/
mkdir fast5/ && cd fast5/
wget -O Klebsiella_pneumoniae_INF032_fast5s.tar.gz \
https://bridges.monash.edu/ndownloader/files/15188573
tar -xzvf Klebsiella_pneumoniae_INF032_fast5s.tar.gz
cd ../
wget -O Klebsiella_pneumoniae_INF032_reference.fasta.gz \
https://bridges.monash.edu/ndownloader/files/14260223
gunzip Klebsiella_pneumoniae_INF032_reference.fasta.gz
```

Cleanup:

```
rm fast5/Klebsiella_pneumoniae_INF032_fast5s.tar.gz
mkdir experiments/
cd ../
```

#### 4.3 *E. coli*

```
mkdir ecolik12mg1655_R10.3/
cd ecolik12mg1655_R10.3/
wget ftp://ftp.sra.ebi.ac.uk/vol1/run/ERR389/ERR3890216/ecolik12mg1655_fast5_R10.3.tar.gz
wget ftp://ftp.ensemblgenomes.org/pub/bacteria/release-47/fasta/bacteria_0_collection/\
escherichia_coli_str_k_12_substr_mg1655/dna/\
Escherichia_coli_str_k_12_substr_mg1655.ASM584v2.dna.toplevel.fa.gz
gunzip Escherichia_coli_str_k_12_substr_mg1655.ASM584v2.dna.toplevel.fa.gz
tar -xzvf ecolik12mg1655_fast5_R10.3.tar.gz
```

Convert to single read fast5 files:

```
multi_to_single_fast5 -i \
srv/MA/nanopore/smk/SMKJ416/LIB-SMKJ416-1-A-1/\
20200130_1036_MN18379_FAL81016_79bb9577/fast5_pass -s fast5
```

Flatten directory structure (source: `https://unix.stackexchange.com/a/52816`):

```
find fast5 -mindepth 2 -type f -exec mv -t fast5 -i '{}' +
```

Cleanup:

```
rm -r ecolik12mg1655_fast5_R10.3.tar.gz srv
mkdir experiments/
cd ../
```

##### 4.4 *H. sapiens*

```
mkdir NA12878 && cd NA12878/
```

Reference:

```
wget ftp://ftp.ensembl.org/pub/release-100/fasta/homo_sapiens/dna/\
Homo_sapiens.GRCh38.dna.primary_assembly.fa.gz
gunzip Homo_sapiens.GRCh38.dna.primary_assembly.fa.gz
```

NA12878 data (single flowcell):

```
mkdir fast5 && cd fast5/
wget http://s3.amazonaws.com/nanopore-human-wgs/rel6/MultiFast5Tars/\
FAB45280-222619780_Multi_Fast5.tar
tar -xvf FAB45280-222619780_Multi_Fast5.tar
multi_to_single_fast5 -i Norwich/FAB45280-222619780_Multi/ -s .
rm -r Norwich/
rm FAB45280-222619780_Multi_Fast5.tar
find . -mindepth 2 -type f -exec mv -t . -i '{}' +
```

Bisulfite data for benchmarking:

```
wget https://www.encodeproject.org/files/ENCFF835NTC/@@download/ENCFF835NTC.bed.gz
gunzip ENCFF835NTC.bed.gz
```

Download NA12878 VCF from GIAB and use bcftools to generate a ground-truth fasta for evaluating basecall accuracy:

```
wget ftp://ftp-trace.ncbi.nlm.nih.gov/ReferenceSamples/giab/release/NA12878_HG001/latest/\
GRCh38/HG001_GRCh38_GIAB_highconf_CG-IllFB-IllGATKHC-Ion-10X-SOLID_CHROM1-X_v.3.3.2_\
highconf_PGandRTGphasetransfer.vcf.gz
wget ftp://ftp-trace.ncbi.nlm.nih.gov/ReferenceSamples/giab/release/NA12878_HG001/latest/\
GRCh38/HG001_GRCh38_GIAB_highconf_CG-IllFB-IllGATKHC-Ion-10X-SOLID_CHROM1-X_v.3.3.2_\
highconf_PGandRTGphasetransfer.vcf.gz.tbi

# handle different chromosome notation in vcf and fasta
for i in {1..22} X Y MT;
do
    echo "chr$i $i" > chr_name_conv.txt
done
$WORKINGDIR/bcftools-1.11/bcftools annotate --rename-chrs chr_name_conv.txt HG001_GRCh38_\
GIAB_highconf_CG-IllFB-IllGATKHC-Ion-10X-SOLID_CHROM1-X_v.3.3.2_highconf_\
```

```
PGandRTGphasetransfer.vcf.gz > \
HG001_GRCh38_GIAB_highconf_CG-IllFB-IllGATKHC-Ion-10X-SOLID_CHROM1-X_v.3.3.2_\
highconf_PGandRTGphasetransfer_nochr.vcf

# compress and index
$WORKINGDIR/bcftools-1.11/bcftools view HG001_GRCh38_GIAB_highconf_CG-IllFB-IllGATKHC-Ion-\
10X-SOLID_CHROM1-X_v.3.3.2_highconf_PGandRTGphasetransfer_nochr.vcf -Oz -o \
HG001_GRCh38_GIAB_highconf_CG-IllFB-IllGATKHC-Ion-10X-SOLID_CHROM_1-X_v.3.3.2_\
highconf_PGandRTGphasetransfer_nochr.vcf.gz
$WORKINGDIR/bcftools-1.11/bcftools index HG001_GRCh38_GIAB_highconf_CG-IllFB-IllGATKHC-Ion-\
10X-SOLID_CHROM1-X_v.3.3.2_highconf_PGandRTGphasetransfer_nochr.vcf.gz

# create NA12878 fasta
$WORKINGDIR/bcftools-1.11/bcftools consensus -p chr -f \
Homo_sapiens.GRCh38.dna.primary_assembly.fa -o NA12878_reference.fa HG001_GRCh38_GIAB_highconf_\
CG-IllFB-IllGATKHC-Ion-10X-SOLID_CHROM1-X_v.3.3.2_highconf_PGandRTGphasetransfer_nochr.vcf.gz
```
